# Experimentally induced fatigue and motor learning: A scoping review

**DOI:** 10.1101/2025.08.07.669206

**Authors:** Bahram Ghafari Goushe, Layale Youssef, Thomas Mangin, Denis Arvisais, Jason L. Neva, Benjamin Pageaux

**Author notes:** equally contributed to this work (Jason L. Neva, Benjamin Pageaux). Corresponding author: Benjamin Pageaux, École de kinésiologie et des sciences de l’activité physique, Centre d’éducation physique et des sports (CEPSUM), Faculté de médecine, Université de Montréal, 2100, boul Édouard-Montpetit, Montreal, Qc, Canada, H3T 1J4.

## Abstract

The literature on the effect of fatigue on motor learning is limited and marked by inconsistent findings. This scoping review aimed to explore the available knowledge on the effects of fatigue induced by physical and cognitive exertion on motor learning, and to compile and understand how it is studied. A comprehensive search strategy using relevant index terms and keywords was conducted across MEDLINE, EMBASE, SPORTDiscus, Web of Science, PsycINFO, CINAHL, ERIC, and Dissertations & Theses Global. Twenty-five studies met the inclusion criteria. The findings revealed considerable inconsistencies in how fatigue and motor learning were defined and measured. None of the studies examined the effect of fatigue induced by combined physical and cognitive exertion, and only 7 studies investigated fatigue induced by cognitive exertion. Acuity tasks were the most frequently used to assess motor learning, employed in 14 studies. Notably, all participants were between 16.5 and 31 years of age, and reporting of key demographic and physiological characteristics such as sex, gender, physical activity level, and body mass index was inconsistent or absent. This review highlights the need for comprehensive definitions of both fatigue and motor learning to improve consistency and reproducibility across studies. Given the limited research on the effects of fatigue induced by cognitive and combined physical and cognitive exertion, future studies should prioritize using these experimental manipulations. Also, future studies should diversify the motor learning tasks used in research to allow both direct and conceptual replication. Additionally, broader age ranges and comprehensive participant profiling should be prioritized.

## 1. Introduction

Skill is defined as the ability to accomplish a task with maximum certainty and minimum energy expenditure (Guthrie, 1935). We have certain innate skills such as walking, running, and balancing that fully develop with maturation and experience (Pringle, 2000). However, to achieve proficiency in other skills that are not innate, we need substantially more practice. Therefore, our lives as human beings are characterized by the performance and learning of skills (Schmidt, 2016; Schmidt, Wrisberg, & Mcdonald, 2006). Since motor skills encompass a large part of human life, for centuries, scientists and educators have been trying to understand the determinants of motor skill learning and the factors that affect their performance (Pringle, 2000).

Motor performance is defined as the observable production of a voluntary motor action at a given moment (Schmidt, 1991). Motor learning is associated with practice and experience that results in long-lasting changes in a person’s capability to respond to task demands (Schmidt, 1991). These changes in capability are inferred from improvements in motor performance. Motor learning has two crucial aspects: 1) skill acquisition – the practice-induced improvement in our ability to rapidly select and then precisely execute appropriate actions; and 2) skill maintenance – the ability to maintain performance levels for a longer period of time beyond skill acquisition and under changing conditions (Krakauer et al., 2019).

Motor learning should be inferred from performance during a delayed retention test that reflects skill maintenance over time. Retention tests allow for the demonstration of maintained performance level, following short– or long-time delays in which the task is not practiced (Magill & Anderson, 2010). Thus, a retention test principally measures performance persistence over time. Another approach to assess motor learning focuses on the generalizability of performance changes. This assessment method involves using transfer tests, which include some novel situation (e.g., novel task parameters), testing the generalizability of the learned task to the characteristics of this new situation (Magill & Anderson, 2010; Pringle, 2000).

Although the most critical requirement to acquire proficiency in a motor skill is practice, fatigue resulting from long bouts of task repetition can be detrimental to motor learning (Branscheidt et al., 2019). Since individuals in the workforce are required to learn and adapt complex motor skills, investigating the interconnection between fatigue and motor learning is of great practical relevance. This is especially critical in fields where mastering intricate fine motor skills is paramount, such as surgery and piloting. Despite a growing interest in the topic of fatigue, many inconsistent findings arise from 1) the lack of a universal definition of fatigue, and 2) the lack of validated experimental designs to induce and measure fatigue (Dong et al., 2022; Enoka & Duchateau, 2016; Mangin & Pageaux, 2025). Fatigue can be induced by physical exertion, cognitive exertion, and a combination of physical and cognitive exertion (Dong et al., 2022; Mangin & Pageaux, 2025). When fatigue is induced by prolonged engagement in physical exertion, it is traditionally defined as muscle or neuromuscular fatigue, which is associated with a reduction in the force production capacity of the working muscles (Gandevia, 2001; Pageaux & Lepers, 2016). When fatigue is induced by prolonged engagement in cognitive exertion, it is traditionally defined as mental or cognitive fatigue, which is associated with a reduction in cognitive performance and/or an increase in subjective feelings of tiredness and lack of energy (Boksem & Tops, 2008; Pageaux & Lepers, 2016; Pageaux & Lepers, 2018). Fatigue is associated with subjective and objective manifestations, and the occurrence of fatigue can be evaluated based on changes in subjective measures (e.g., increases in self-reported fatigue or in the perceived effort required to maintain performance in its presence) and in objective physiological and behavioral measures (e.g., increases in heart rate with a constant workload or increases in reaction time) (Dong et al., 2022; Mangin & Pageaux, 2025).

To the best of our knowledge, few studies have investigated the effect of fatigue induced by physical exertion on motor learning, and results are inconsistent (Alderman, 1965; Carron, 1969; Cotten et al., 1972). Regarding fatigue induced by cognitive exertion, to the best of our knowledge, only one study has been published, and it suggests a positive effect of fatigue (Borragan et al., 2016). There is also a large variability in the methods used to infer the presence of fatigue that may contribute to the heterogeneity of the results observed in the literature. Therefore, a systematic exploration of the methodologies and findings of the relevant literature is needed for a better understanding of the effects of fatigue on motor learning.

In this context, this scoping review aimed to explore the available knowledge on the effects of fatigue induced by physical and cognitive exertion on motor learning. It synthesized the key concepts and definitions in the fields of fatigue and motor learning, examined the methodologies used to induce and measure fatigue, and to infer motor learning. Finally, this scoping review identified gaps in the existing literature and proposed recommendations for future research. More specifically, this review sought to investigate (1) the methods used to induce and quantify fatigue, (2) the methods used to assess motor learning, (3) how fatigue and motor learning were defined, and (4) the target populations that had been examined.

## 2. Methods

The protocol was pre-registered in the Open Science Framework (https://osf.io/39m2a/resources) and was conducted according to the JBI scoping review guidelines and recommendations (Peters et al., 2020). Any deviation from the pre-registration is reported.

### 2.1 Search strategy and data extraction

A three-step search strategy was used. A preliminary search of OSF, MEDLINE, Cochrane Database of Systematic Reviews, and the JBI Evidence Synthesis conducted in September 2021 yielded no completed or ongoing scoping and systematic reviews on the topic. The text words contained in the titles and abstracts of relevant articles, as well as the index terms used to describe the papers, were analyzed. After that, a second search was conducted in collaboration with a librarian (DA) in November 2021. A final search was conducted by DA in June 2025. For the concept of fatigue, included terms were “mental fatigue,” “physical fatigue,” “cognitive fatigue,” “cognitive exertion,” and “physical exertion.” For the concept of motor learning, terms such as “motor learning,” “motor skill,” “motor adaptation,” “motor acuity,” “motor sequence”, and “motor acquisition” were included. All the identified index terms and keywords were used to search across the databases, including MEDLINE (1946-), EMBASE (1974-), PsychINFO (1806), SPORTDiscus (1975-), CINAHL (1937-), Web of Science (1945-), ProQuest (1966), and Dissertations & Theses Global. The detailed search equations are provided in supplementary material 1.

#### 2.1.1 Screening and selection of sources of evidence

All identified articles were uploaded to the screening and data extraction tool Covidence. Duplicates were removed, and titles and abstracts were screened according to the review inclusion criteria by two independent researchers, BGG and LY. Full texts of the remaining studies were retrieved and examined according to the review inclusion criteria by the same two independent researchers. Studies that did not meet the inclusion criteria were excluded, and reasons for the exclusion were provided and reported in the flowchart. In case the two reviewers failed to reach a consensus, JN and BP were brought in to resolve the disagreement. A detailed flowchart of the inclusion process is available in Figure 1.

**Figure 1.**
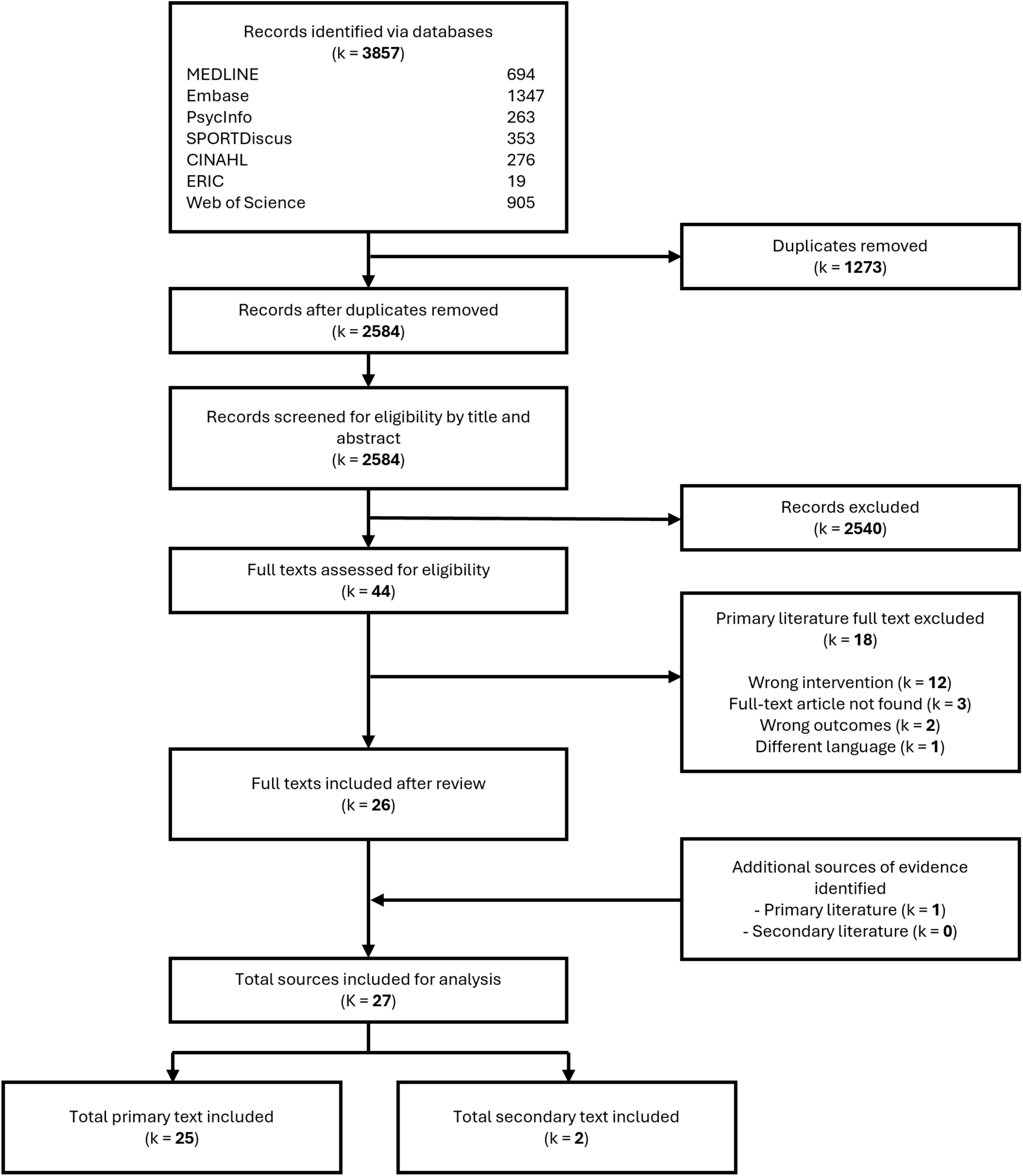
PRISMA flow chart for the identification, screening, and inclusion of articles. k represents the number of texts.

#### 2.1.2 Data extraction

Data extraction was conducted using a data extraction table. The data extraction table presented in the pre-registration was updated to capture more detailed information about the articles. A preliminary data extraction table was piloted on the first three studies by the authors (BGG, LY, JN, and BP). The finalized data extraction table, available in supplementary material 2 was used by BGG and LY to extract the information from each full-text article. Disagreements were resolved through discussion.

### 2.2 Inclusion criteria

#### 2.2.1 Sources of evidence

All types of fully published peer-reviewed articles (systematic reviews, quantitative, qualitative, conference proceedings, theses, and dissertations), as well as the gray literature, were eligible for inclusion. The language was restricted to English and French.

#### 2.2.2 Population

This scoping review included studies that involved healthy participants of any gender and sex. This scoping review focused on fatigue induced by voluntary engagement in a task and not fatigue resulting from pathological conditions (e.g., chronic fatigue syndrome, stroke, etc.). This choice was made to control for the confounding effects of the pathologies on fatigue.

#### 2.2.3 Motor learning

Motor learning is associated with practice and experience that results in long-lasting changes in a person’s capability to respond to task demands (Schmidt, 1991). Studies were included if they required participants to practice a motor task and/or incorporated either a retention or transfer test to confirm that motor learning had occurred. To classify the motor learning tasks used across the included studies, we identified four main categories based on task characteristics and learning objectives (e.g., Krakauer et al., 2019). This classification is used in a scoping review of our research group on the effect of physical exercise on motor learning (Youssef et al., 2025). Sequence learning, often referred to as sequence-specific learning, was defined as motor tasks that involve comparing performance on repeated sequences with that on random sequences (e.g., serial reaction time task; Robertson, 2007). This comparison is essential to identify learning that is specific to the practiced sequence, distinguishing it from general improvements in other aspects of the motor skill. However, when only repeated sequences were practiced without a random condition for comparison, the task was classified as motor acuity. Motor acuity refers to improvements in the accuracy and/or precision of performing a particular movement or sequence of movements through practice (e.g., Rho task; Alderman, 1965), with a focus on refining the ability to execute the motor task more precisely and/or accurately. Motor adaptation included tasks in which participants adjusted their movements in response to external perturbation (e.g., visuomotor rotation task; Neva et al., 2019). Associative learning included tasks requiring participants to form arbitrary associations between stimuli and responses (e.g., conditional learning task; Balsters & Ramnani, 2011). The primary objective of each task category is shown in Figure 2. Although these task categories were not always mutually exclusive, for the sake of this review, we classified each study into a single category. In cases where categorization conflicts arose, the authors resolved them through discussion and consensus. As a deviation from the pre-registration, the classification of motor learning task categories has been added to enable us to identify the gaps in the methods used to study motor learning.

**Figure 2.**
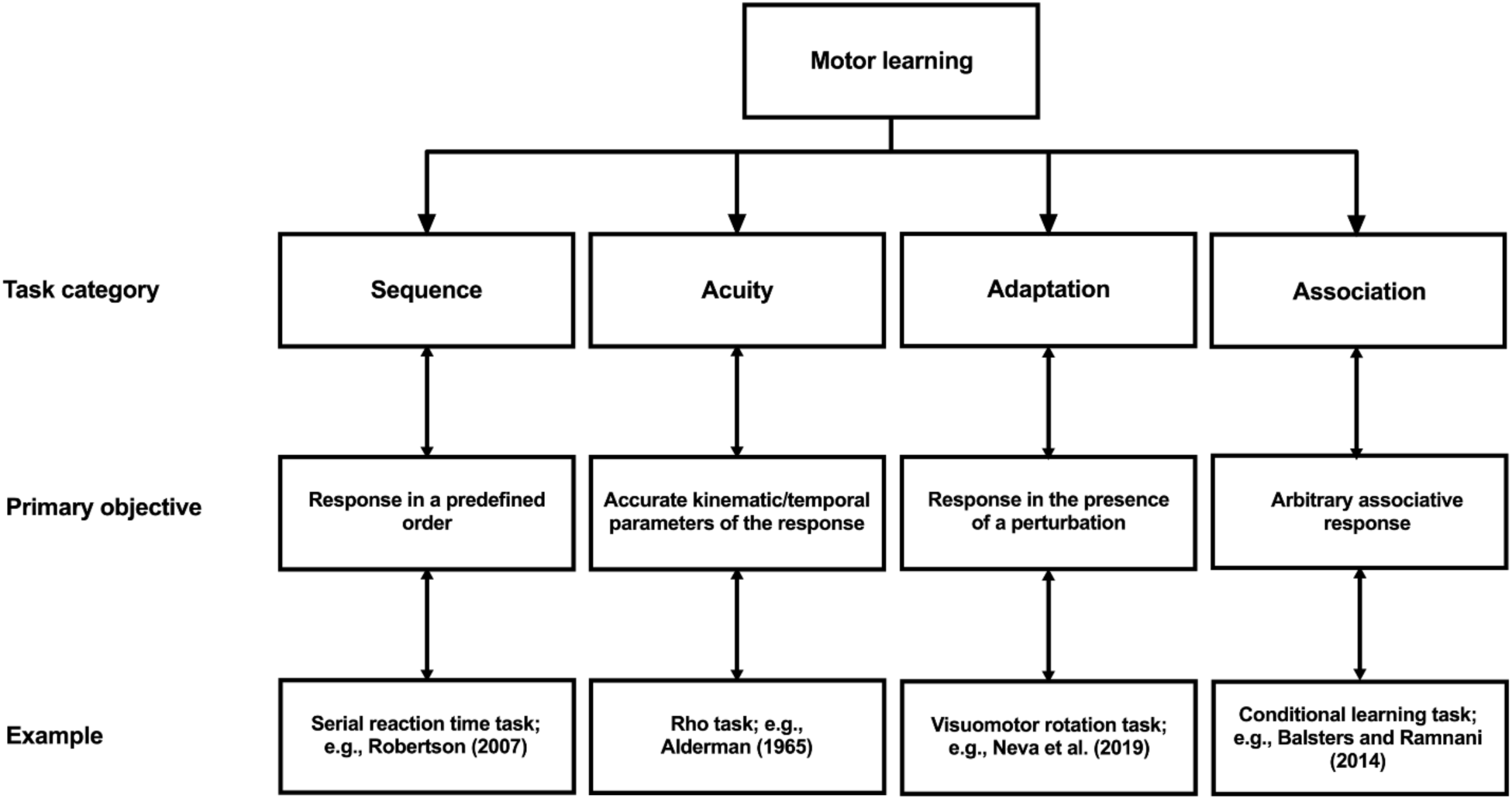
Classification of the task categories investigated to infer motor learning. This figure illustrates four commonly used motor learning task categories, classified according to their primary objective. The examples shown represent typical experimental tasks used within each category to assess motor learning. The figure is reproduced with permission from the supplementary material of a scoping review of our research group (preprint license CC-BY-NC-ND 4.0) on the effect of physical exercise on motor learning (Youssef et al., 2025).

#### 2.2.4 Fatigue

Fatigue is a psychobiological phenomenon induced by prolonged engagement in physical and/or cognitive exertion (Behrens et al., 2023; Mangin & Pageaux, 2025; Pageaux & Lepers, 2016). The presence of fatigue had to be confirmed by measurement of subjective and/or objective manifestations of fatigue. Subjectively, fatigue manifests as increased feelings of tiredness or exhaustion (e.g., increases in self-reported fatigue over time). Objectively, if the level of engagement in a task remains constant, fatigue impairs physical and cognitive performance (e.g., behavioral: increase in reaction time; or physiological: increased heart rate with the same workload). The included studies had to apply a fatiguing protocol performed during the same session as the motor learning task and include at least one criterion to verify that fatigue had been successfully induced (i.e., manipulation check to confirm fatigue induction). Figure 3 shows the classification used to describe the fatigue induced. As a deviation from the pre-registration, the classification of combined physical and mental fatigue has been added to enable us to identify the gaps in the methods used to study fatigue. Moreover, self-reported measures of effort were considered, as effort is related to fatigue in such a way that, in the presence of fatigue, performance can be maintained through compensatory mechanisms involving an increased investment of effort in the task (Wright & Mlynski, 2019). Incorporating self-reported measures of effort is also an update to the pre-registration.

**Figure 3.**
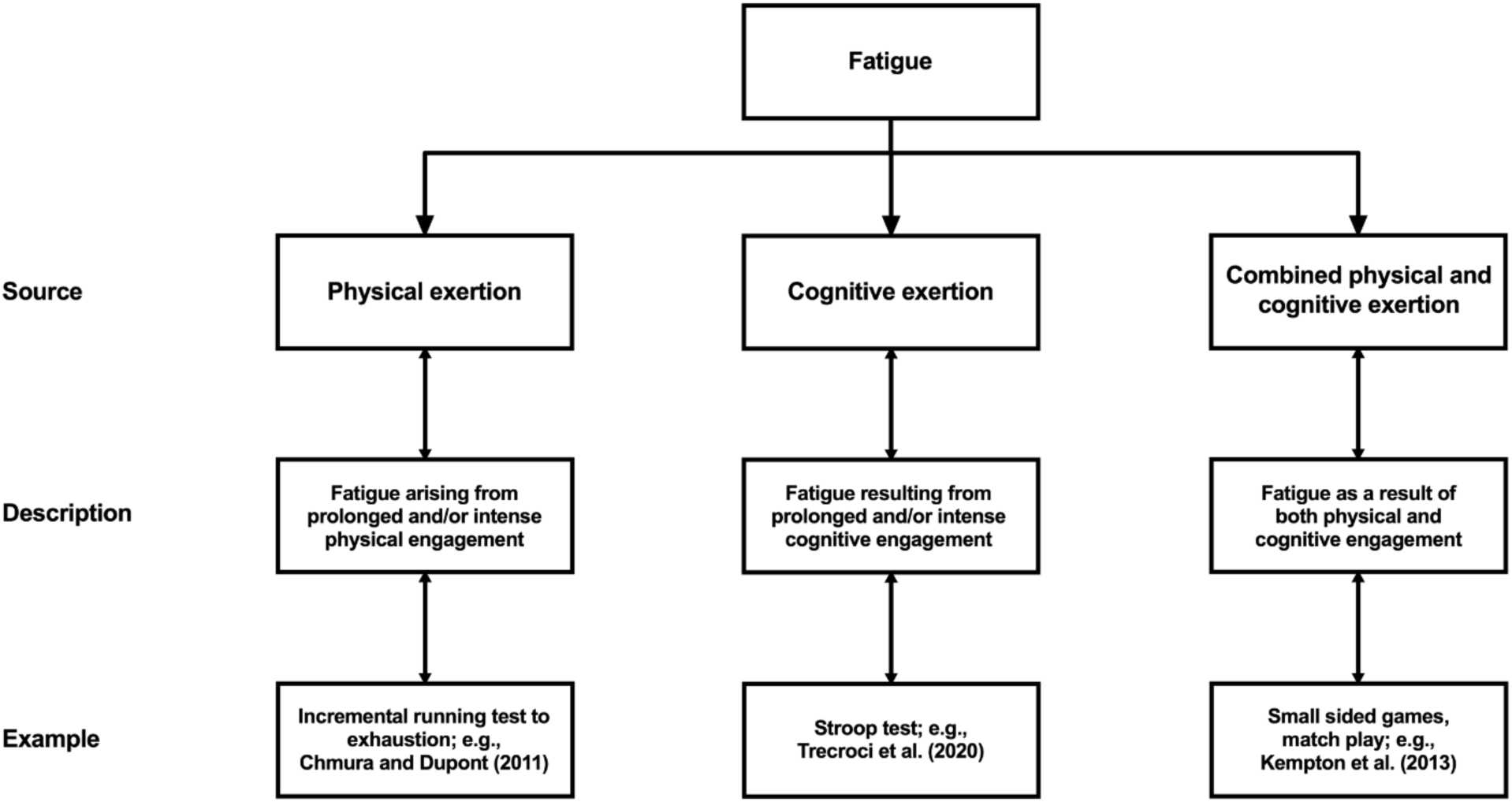
Classifications used to describe the fatigue induced. This figure illustrates three sources of fatigue along with corresponding descriptions. The examples provided represent typical experimental tasks used in the literature to induce fatigue.

## 3. Results

### 3.1 Studies included

The flow chart in Figure 1 presents the results of the search along with reasons for exclusion. The initial database search returned 2709 references, of which 36 full texts were screened for inclusion. An additional article was identified through forward citation searching. The updated search in June 2025 returned 1142 references, of which 3 full texts were included. Following peer review, three additional articles were suggested for consideration, of which one met the inclusion criteria and was included in the review. Reviewers also recommended adding the term *“*motor acquisition*”* to our original search strategy. This updated search, conducted in November 2025, identified two additional articles; one met the inclusion criteria and was included. In addition, another study was published after the updated search and was identified through Google Scholar citation update received by one of the authors. As this study fulfilled the inclusion criteria, it was also included. As a result, a total of 3 additional articles were included following the peer-review process.

To ensure transparency and reproducibility in the full-text screening process, the reasons for exclusion of the 18 screened studies are reported in Supplementary Material 3 (Table S1)

In total, 27 sources were included: 25 primary sources (original research articles) and 2 secondary sources (1 MSc thesis and 1 book chapter). These 27 sources were categorized for analysis as 25 experimental studies, 1 MSc thesis, and 1 book chapter.

A timeline of published articles is presented in Figure 4A. The topic of fatigue’s effect on motor learning was initially studied in the 1960s with 3 studies (Alderman, 1965; Carron, 1969; Nunney, 1963). This research topic received more attention in the 1970s, with 5 studies (Cotten et al., 1974; Cotten et al., 1972; Dickinson, Medhurst, & Whittingham, 1979; Godwin & Schmidt, 1971; Whitley, 1975), then slowed down, with only one study published almost every decade (Arnett, DeLuccia, & Gilmartin, 2000; Berger & Smith-Hale, 1991; Dwyer, 1984) until the early 2000s (Masters, Poolton, & Maxwell, 2008; Mierau et al., 2009; Takahashi et al., 2006). Interest in the topic increased again between 2011 and 2025, with 11 studies (Anguera et al., 2012; Apreutesei & Cressman, 2024; Banihosseini, Abdoli, & Kavyani, 2025; Borragan et al., 2016; Branscheidt et al., 2019; Godoi Filho et al., 2025; Hoskens et al., 2022; Khojasteh Moghani et al., 2021; Nardon et al., 2024; Siekirk, Lai, & Kendall, 2019; Zabihhosseinian et al., 2020).

**Figure 4.**
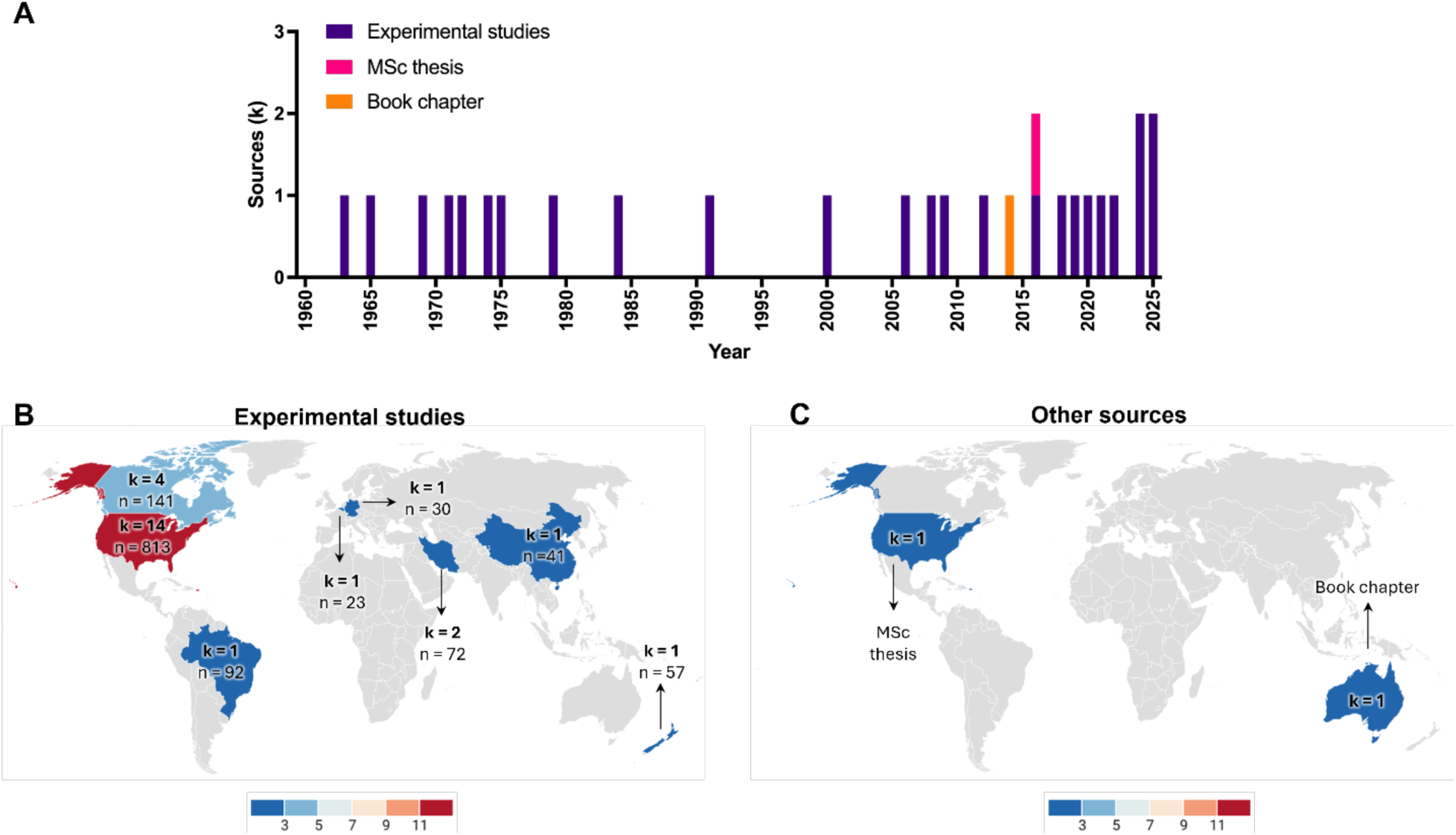
(A) Timeline of studies investigating the effect of fatigue on motor learning from the 1960s to 2025. (B) Geographical distribution of the experimental studies included (k), with the corresponding number of participants (n). (C) Geographical distribution of the other sources included. One MSc thesis from the United States of America and one book chapter from Australia.

All 25 articles were in English. The origin of the study is defined based on the corresponding author information. Included articles originated from the United States of America (k = 14), Canada (k = 4), Iran (k = 2), Belgium (k = 1), New Zealand (k = 1), China (k = 1), Brazil (k = 1), and Germany (k = 1). The geographical origin of the included studies is presented in Figure 4B. All the articles were laboratory studies. Twenty-three articles (92%) included both within and between-subject designs, and two included a within-subject design (8%). For the 2 secondary sources, the MSc thesis (Datla, 2016) originated from the United States of America, and the book chapter (Taylor, 2013) originated from Australia (Figure 4C).

We extracted data from 25 experimental studies to address our specific research questions. The following sections address our review questions. An overview of the main characteristics of all experimental studies is presented in Table 1. Detailed characteristics of all experimental studies are provided in Table S2 of the supplementary material 4.

**Table 1.**
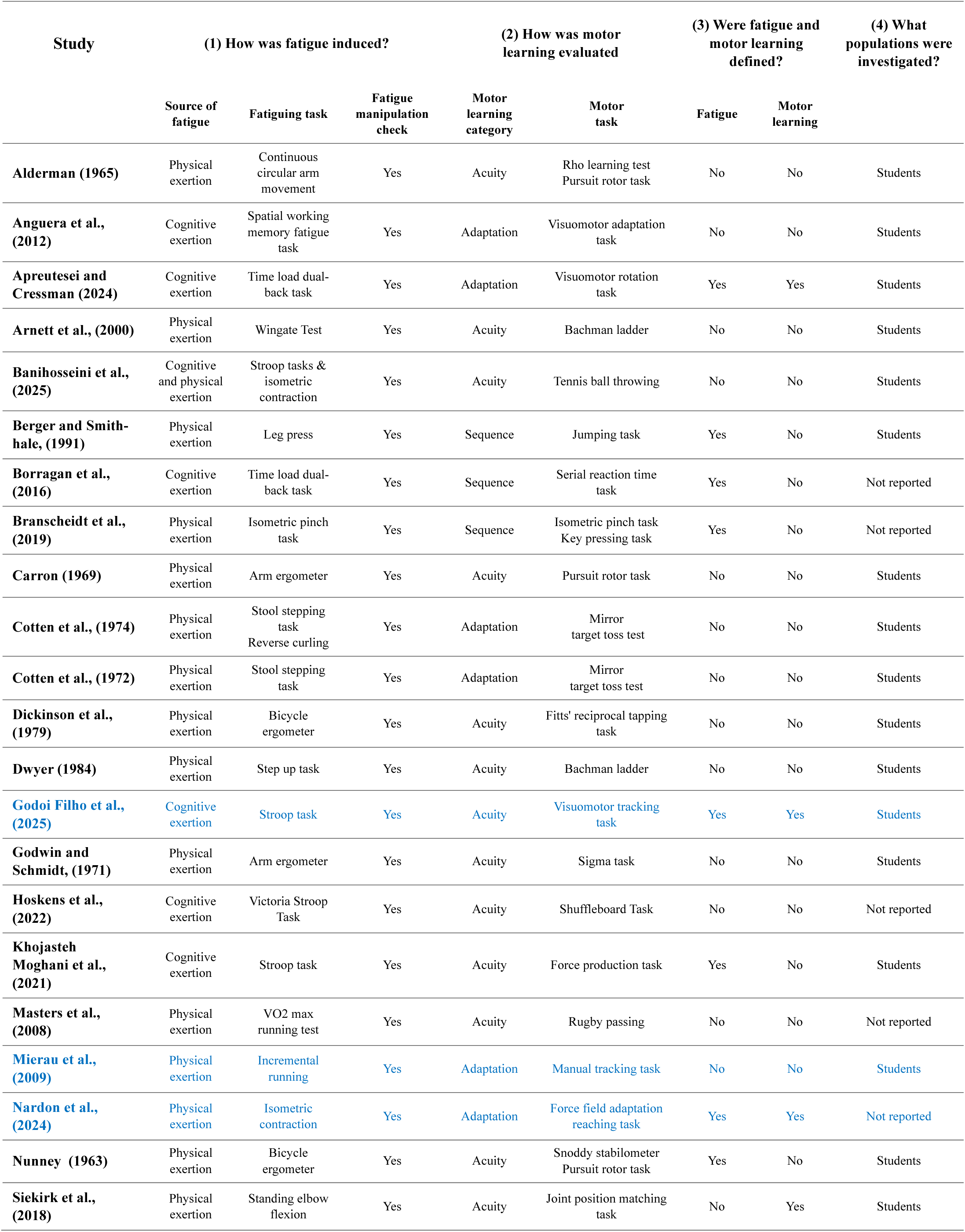
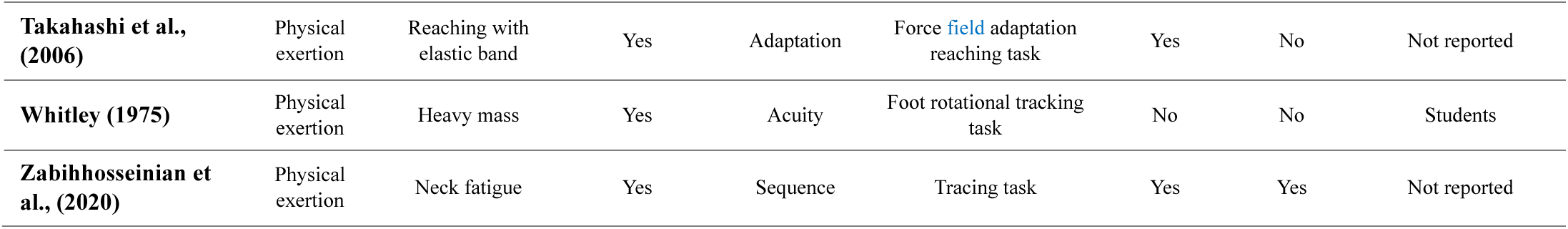
Characteristics of the included studies related to the review questions.

### 3.2 Methods to induce and measure fatigue

#### 3.2.1 Source of fatigue

Figure 5A shows the data on sources of fatigue. One study involved two tasks to induce physical fatigue, stool stepping task and reverse curling task (Cotten et al., 1974), and one study (Banihosseini, Abdoli, & Kavyani, 2025) used both mental and physical exertion to induce fatigue, employing a Stroop task and an isometric contraction task, respectively. In 27% of studies (k = 7), fatigue was induced by cognitive exertion. The cognitive exertion involved spatial working memory task (Anguera et al., 2012), time load dual-back task (Apreutesei & Cressman, 2024; Borragan et al., 2016), and Stroop task (Banihosseini, Abdoli, & Kavyani, 2025; Godoi Filho et al., 2025; Hoskens et al., 2022; Khojasteh Moghani et al., 2021). In 73% of the studies (k =19), fatigue was induced by physical exertion. The physical exertion included the use of an arm ergometer (Carron, 1969; Godwin & Schmidt, 1971), a bicycle ergometer (Dickinson, Medhurst, & Whittingham, 1979; Nunney, 1963), continuous circular arm movements (Alderman, 1965), moving a heavy mass (Whitley, 1975), an isometric pinch task (Branscheidt et al., 2019), leg press (Berger & Smith-Hale, 1991), neck contractions (Zabihhosseinian et al., 2020), reaching with an elastic band (Takahashi et al., 2006), standing elbow flexion (Siekirk, Lai, & Kendall, 2019), a step up task (Dwyer, 1984), a V̇O_2max_ running test (Masters, Poolton, & Maxwell, 2008), a Wingate test (Arnett, DeLuccia, & Gilmartin, 2000), a stool stepping task (Cotten et al., 1974; Cotten et al., 1972), reverse biceps curls (Cotten et al., 1974), an isometric contraction task (Banihosseini, Abdoli, & Kavyani, 2025; Nardon et al., 2024), and an incremental running task (Mierau et al., 2009). No study employed a combined cognitive and physical exertion method to induce fatigue.

**Figure 5.**
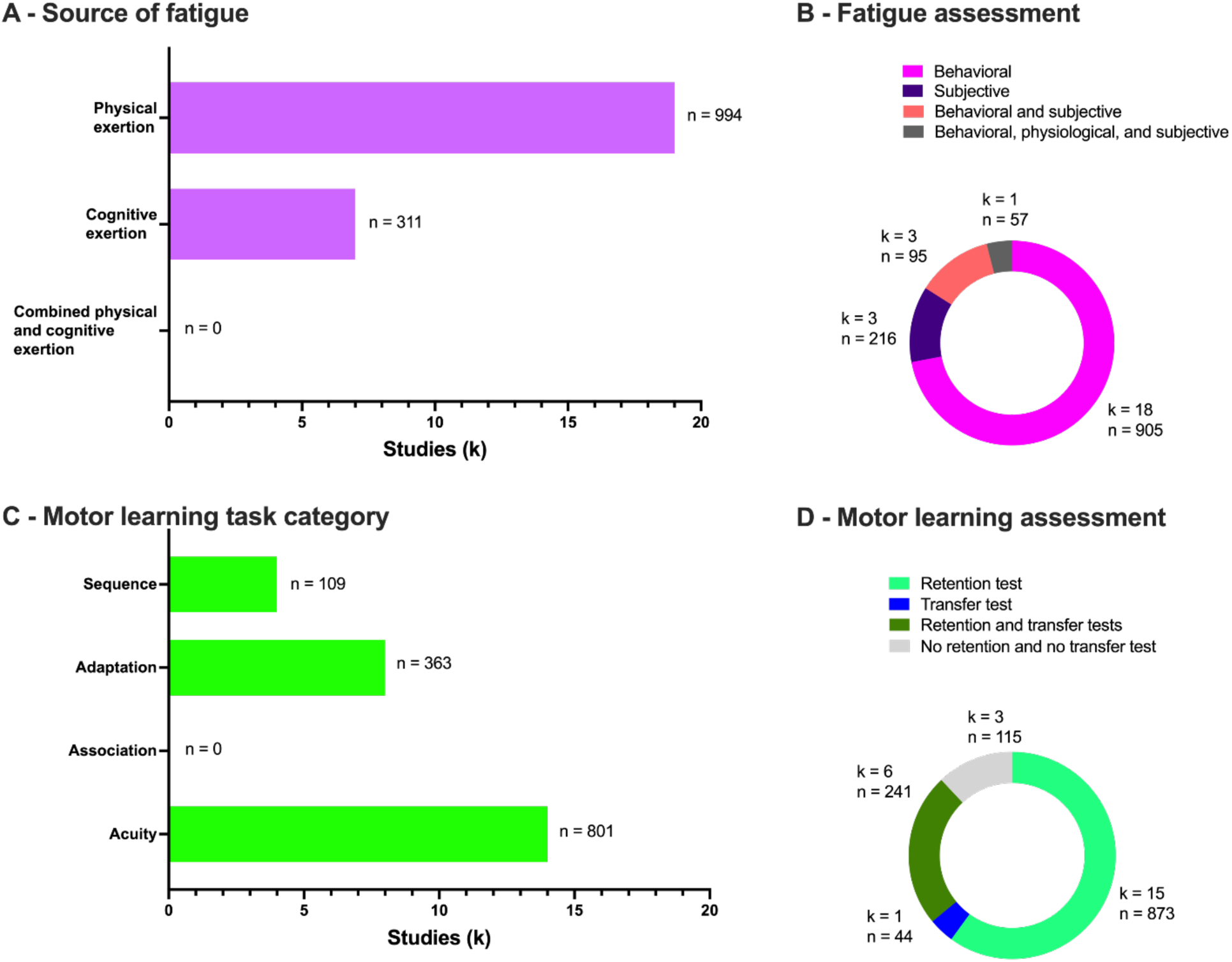
(A) Number of studies (k) that used physical tasks only, cognitive tasks only, or a combination of both, with the corresponding number of participants (n). (B) Number of studies that used objective (behavioral or physiological), subjective, and a combination of both objective (behavioral or physiological) and subjective manipulation checks to confirm the successful induction of fatigue. (C) Number of studies that investigated a sequence, adaptation, association, or acuity task category, with the corresponding number of participants. (D) Number of studies that used retention test, transfer test, and a combination of both retention and transfer tests to confirm learning has occurred.

#### 3.2.2 Criterion to verify the successful fatigue induction

Figure 5B shows the data on the criterion to verify the successful fatigue induction. Of all the designs of the 25 included studies, 12% of the studies (k = 3) attested to the presence of fatigue by exclusively measuring its subjective manifestation. These studies measured subjective feelings of fatigue using a multidimensional fatigue inventory (Khojasteh Moghani et al., 2021), a nine-point subjective rating scale (Nunney, 1963), and a visual analog scale (Godoi Filho et al., 2025). Seventy-two percent of the studies (k = 18) verified the presence of fatigue by exclusively measuring its objective manifestation. All these studies measured changes in task performance, i.e., behavioral manipulation check (Alderman, 1965; Anguera et al., 2012; Arnett, DeLuccia, & Gilmartin, 2000; Berger & Smith-Hale, 1991; Branscheidt et al., 2019; Nardon et al., 2024; Whitley, 1975; Zabihhosseinian et al., 2020), and volitional exhaustion tests (Carron, 1969; Cotten et al., 1974; Cotten et al., 1972; Dickinson, Medhurst, & Whittingham, 1979; Dwyer, 1984; Godwin & Schmidt, 1971; Masters, Poolton, & Maxwell, 2008; Mierau et al., 2009; Siekirk, Lai, & Kendall, 2019; Takahashi et al., 2006). Twelve percent (k = 3) of the included studies confirmed the presence of fatigue by measuring both objective (behavioral) and subjective manifestations (Apreutesei & Cressman, 2024; Banihosseini, Abdoli, & Kavyani, 2025; Borragan et al., 2016). One study (Hoskens et al., 2022) employed a physiological measurement (electroencephalography; EEG T7-Fz connectivity) in addition to behavioral and subjective measurements to further demonstrate the presence of fatigue. No study utilized a self-reported measure of effort.

### 3.3 Methods to assess motor learning

#### 3.3.1 Task categories investigated to infer motor learning

Figure 5C shows the data on task categories to infer motor learning. Two studies used more than one task to assess motor learning. One of these studies used two sequence tasks (Branscheidt et al., 2019) and the other one used the acuity and adaptation tasks (Nunney, 1963). Sixteen percent of the studies (k = 4) employed tasks falling within the sequence task category. The sequence task category involved the use of the serial reaction time task (Borragan et al., 2016), isometric pinch task (Branscheidt et al., 2019), key-pressing task (Branscheidt et al., 2019), tracing task (Zabihhosseinian et al., 2020), and jumping task (Berger & Smith-Hale, 1991). Twenty-eight percent of the articles (k = 7) employed a sensorimotor adaptation task. The studies investigating the sensorimotor adaptation category included the visuomotor adaptation task (Anguera et al., 2012), mirror target toss test (Cotten et al., 1974; Cotten et al., 1972), Snoddy stabilometer (Nunney, 1963), force-field adaptation reaching task (Nardon et al., 2024; Takahashi et al., 2006), visuomotor rotation task (Apreutesei & Cressman, 2024), and manual tracking task (Mierau et al., 2009). Fifty-six percent of the articles (k = 14) adopted tasks that fell into the acuity task category. Within the acuity task category, tasks included the Rho learning test (Alderman, 1965), pursuit rotor task (Alderman, 1965; Carron, 1969; Nunney, 1963), Bachman ladder (Arnett, DeLuccia, & Gilmartin, 2000; Dwyer, 1984), Fitts’ reciprocal tapping task (Dickinson, Medhurst, & Whittingham, 1979), Sigma task (Godwin & Schmidt, 1971), force production task (Khojasteh Moghani et al., 2021), rugby passing skill (Masters, Poolton, & Maxwell, 2008), foot rotational tracking task (Whitley, 1975), joint-position matching task (Siekirk, Lai, & Kendall, 2019), Tennis ball throwing task (Banihosseini, Abdoli, & Kavyani, 2025), Shuffleboard task (Hoskens et al., 2022), and visuomotor tracking task (Godoi Filho et al., 2025).

The time interval between the fatiguing task and motor learning task was reported in 16% of the studies (k = 4). This time interval ranged from 3-5 seconds (Arnett, DeLuccia, & Gilmartin, 2000; Carron, 1969; Godwin & Schmidt, 1971) to 45 seconds (Masters, Poolton, & Maxwell, 2008). All 4 studies involved fatigue induced by physical exertion.

#### 3.3.2 Motor learning assessment

Figure 5D shows the data on motor learning assessment methods. Sixty percent of the studies (k = 15) exclusively employed a retention test to confirm the occurrence of motor learning (Alderman, 1965; Anguera et al., 2012; Apreutesei & Cressman, 2024; Berger & Smith-Hale, 1991; Borragan et al., 2016; Carron, 1969; Cotten et al., 1974; Cotten et al., 1972; Dickinson, Medhurst, & Whittingham, 1979; Dwyer, 1984; Godoi Filho et al., 2025; Nunney, 1963; Takahashi et al., 2006; Whitley, 1975; Zabihhosseinian et al., 2020). One study used a transfer test exclusively to infer motor learning (Arnett, DeLuccia, & Gilmartin, 2000). Twenty-seven percent of the articles (k = 6) used a transfer test in addition to a retention test (Banihosseini, Abdoli, & Kavyani, 2025; Branscheidt et al., 2019; Godwin & Schmidt, 1971; Khojasteh Moghani et al., 2021; Masters, Poolton, & Maxwell, 2008; Siekirk, Lai, & Kendall, 2019). Twelve percent of studies (k = 3) measured motor acquisition and did not include any retention or transfer tests (Hoskens et al., 2022; Mierau et al., 2009; Nardon et al., 2024).

### 3.4 How are fatigue and motor learning defined?

Table 2 reports whether a definition of fatigue and motor learning was provided for each included study, as well as the definition reported. Among all the reviewed articles, 16% (k = 4) included both the definitions of fatigue and motor learning (Apreutesei & Cressman, 2024; Godoi Filho et al., 2025; Nardon et al., 2024; Zabihhosseinian et al., 2020). However, the definition given for fatigue by Zabihhosseinian et al., (2020) describes the concept of fatigability, i.e., specific information related to the declined performance component of fatigue (Kluger, Krupp, & Enoka, 2013; Skau, Sundberg, & Kuhn, 2021) rather than defining fatigue. Twenty-four percent of the studies (k = 6) only included a definition of fatigue (Berger & Smith-Hale, 1991; Borragan et al., 2016; Branscheidt et al., 2019; Khojasteh Moghani et al., 2021; Nunney, 1963; Takahashi et al., 2006), and one article only included a definition of motor learning (Siekirk, Lai, & Kendall, 2019). Fifty-six percent of the articles (k = 14) included neither a definition of fatigue nor motor learning (Alderman, 1965; Anguera et al., 2012; Arnett, DeLuccia, & Gilmartin, 2000; Banihosseini, Abdoli, & Kavyani, 2025; Carron, 1969; Cotten et al., 1974; Cotten et al., 1972; Dickinson, Medhurst, & Whittingham, 1979; Dwyer, 1984; Godwin & Schmidt, 1971; Hoskens et al., 2022; Masters, Poolton, & Maxwell, 2008; Mierau et al., 2009; Whitley, 1975). While 56% (k = 14) of articles did not include an explicit definition of fatigue, it is nonetheless possible to use some information provided by the authors to infer the fatigue framework used in each study. Across all included articles, 8% (k = 2) did not provide sufficient information to determine the fatigue framework used in their study. The remaining 92% (k = 24) of articles could be classified into three main conceptual frameworks, as reported in Table S3 of the supplementary material 5. In 38% (k = 9) of these articles, fatigue is conceptualized as an objective performance decrement, and 17% (k = 4) specific to motor performance only. In 8% (k = 2) of articles, fatigue is conceptualized as a depletion of cognitive resources. In 8% (k = 2) articles, fatigue is conceptualized as subjective feelings of tiredness or lack of energy. Twenty-one percent (k = 5) combined the objective performance decrement and the depletion of cognitive resources. Finally, 8% (k = 2) of articles combined the objective performance decrement with the subjective feelings of tiredness or lack of energy framework.

**Table 2.**
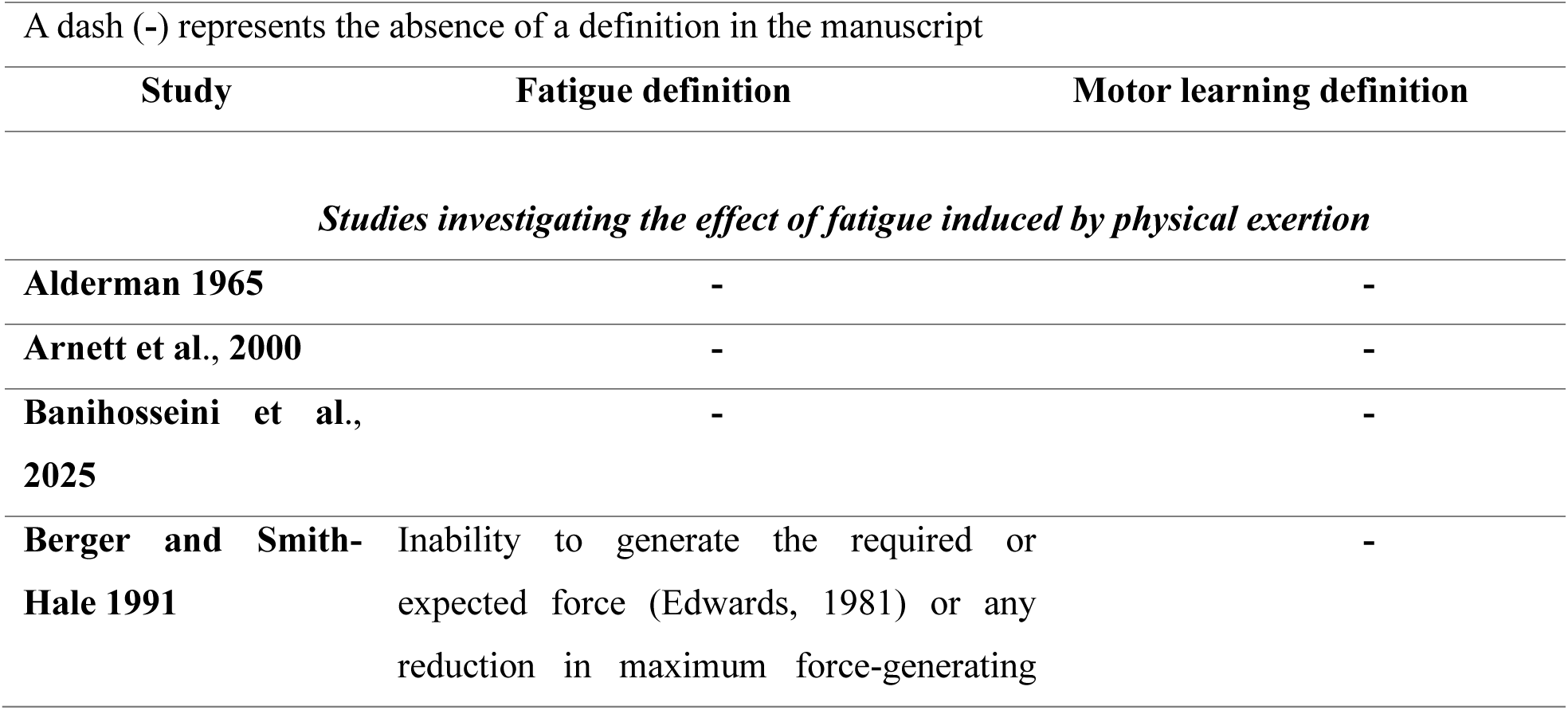
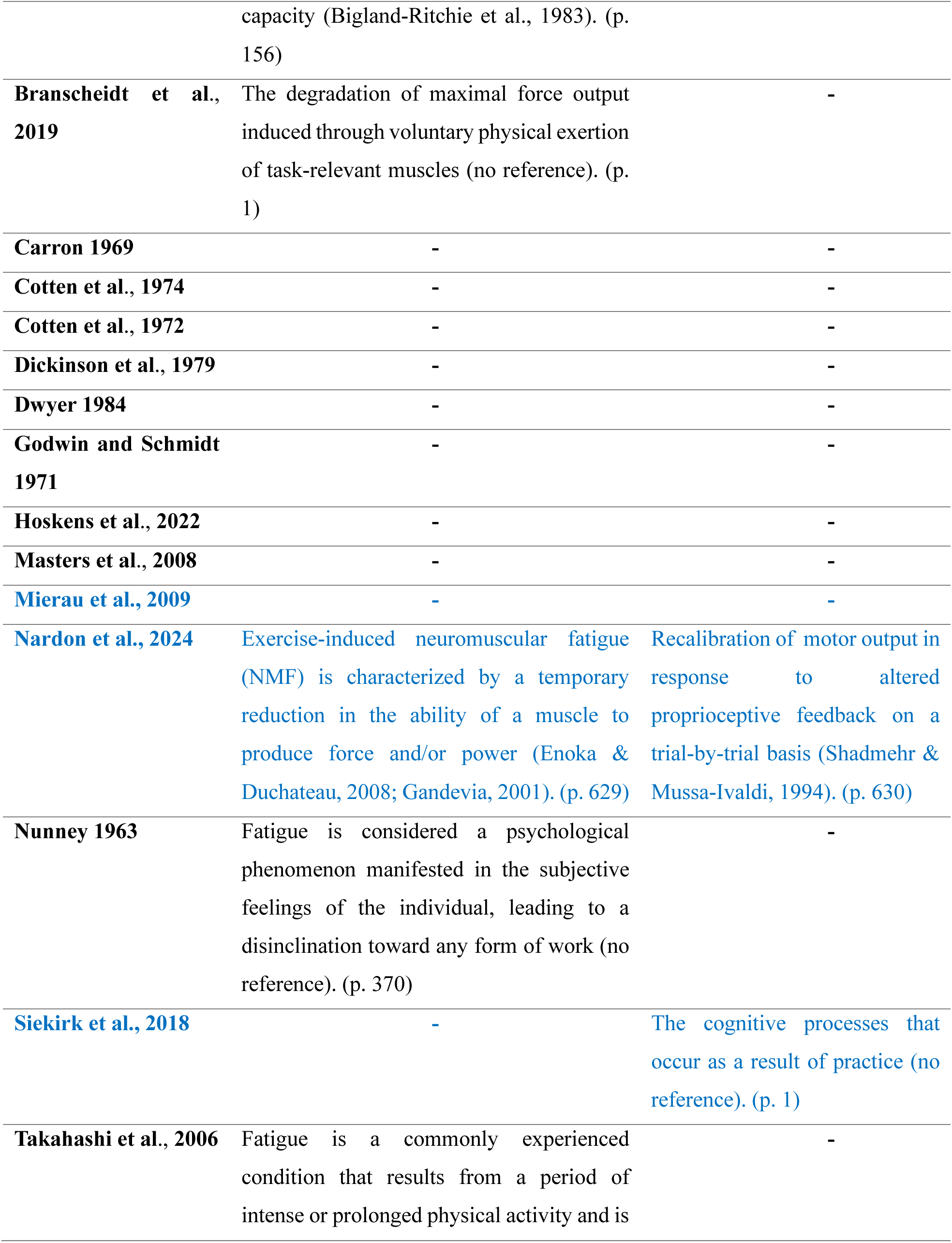
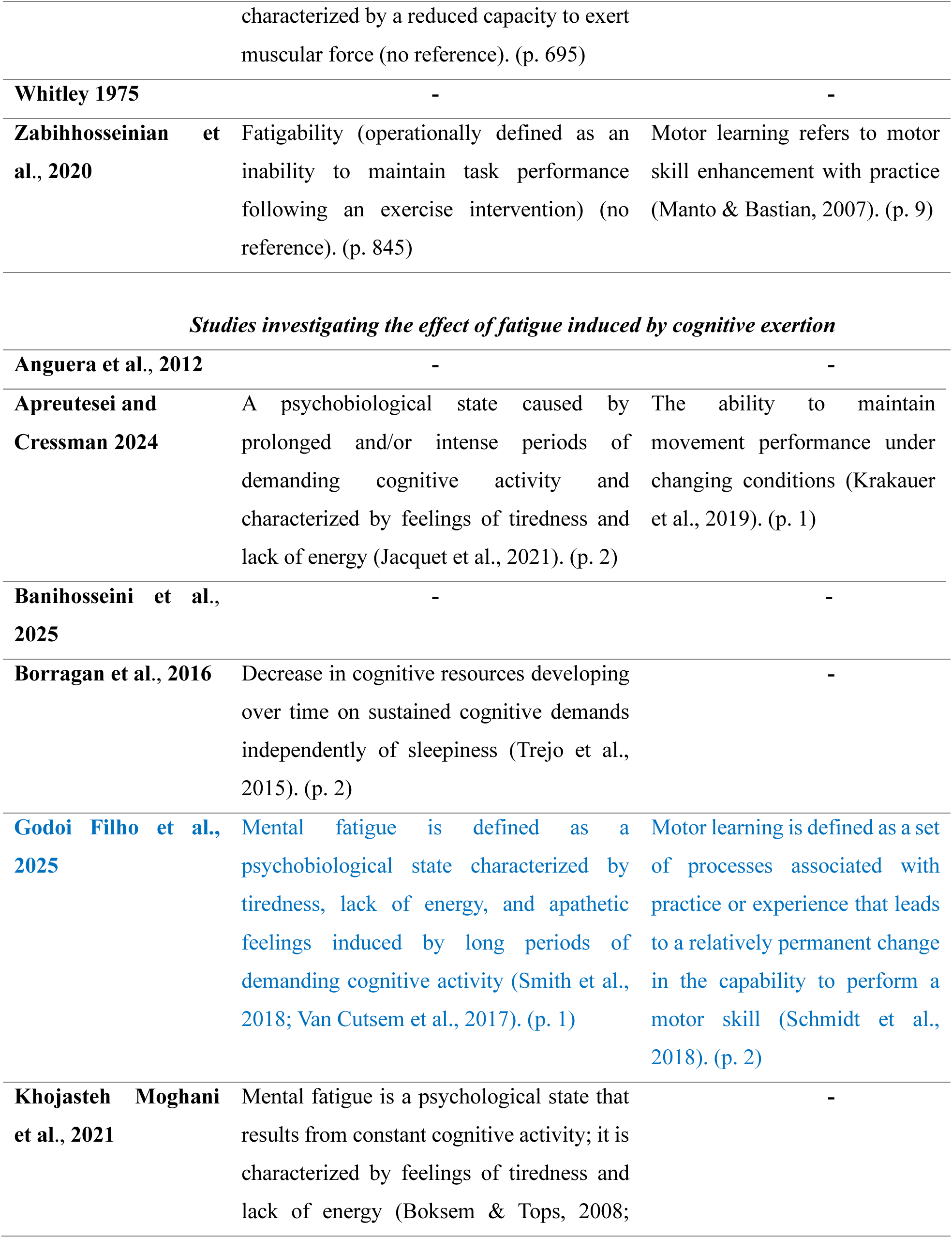
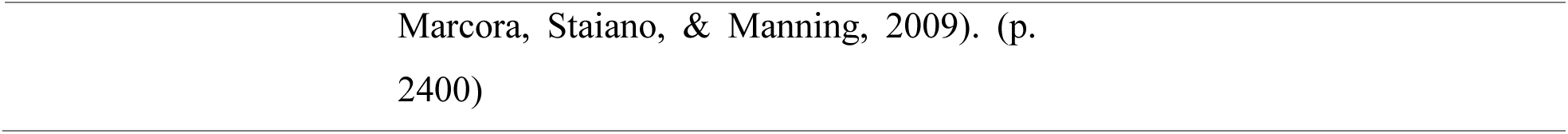
Studies that included a definition for fatigue, motor learning, or both.

Overall, the definitions of fatigue and motor learning were inconsistent across studies. The definitions provided for fatigue are confined within a specific discipline rather than encompassing fatigue as a multidomain phenomenon. Similarly, the two definitions of motor learning lacked a core element, i.e., its characterization as a process leading to long-lasting improvements in performance (Table 2).

### 3.5 Participants

Figure 6 shows the data on participant characteristics. Twenty-five studies with 35 participant groups were considered. Regarding sex, 36% of the studies (k = 9) exclusively involved male participants (Alderman, 1965; Berger & Smith-Hale, 1991; Cotten et al., 1974; Cotten et al., 1972; Dwyer, 1984; Khojasteh Moghani et al., 2021; Mierau et al., 2009; Nunney, 1963; Whitley, 1975). Sixteen percent of studies (k = 4) recruited both male and female participants (Apreutesei & Cressman, 2024; Dickinson, Medhurst, & Whittingham, 1979; Hoskens et al., 2022; Nardon et al., 2024). Forty-eight percent of studies (k = 12) did not report any information about the participants’ sex (Anguera et al., 2012; Arnett, DeLuccia, & Gilmartin, 2000; Banihosseini, Abdoli, & Kavyani, 2025; Borragan et al., 2016; Branscheidt et al., 2019; Carron, 1969; Godoi Filho et al., 2025; Godwin & Schmidt, 1971; Masters, Poolton, & Maxwell, 2008; Siekirk, Lai, & Kendall, 2019; Takahashi et al., 2006; Zabihhosseinian et al., 2020) (Figure 6A&B). Although we categorized the available data under “sex” or “gender” based on the original reporting in each article, we acknowledge that these terms are often used interchangeably in the literature, despite referring to distinct concepts (e.g., Diamond, 2002).

**Figure 6.**
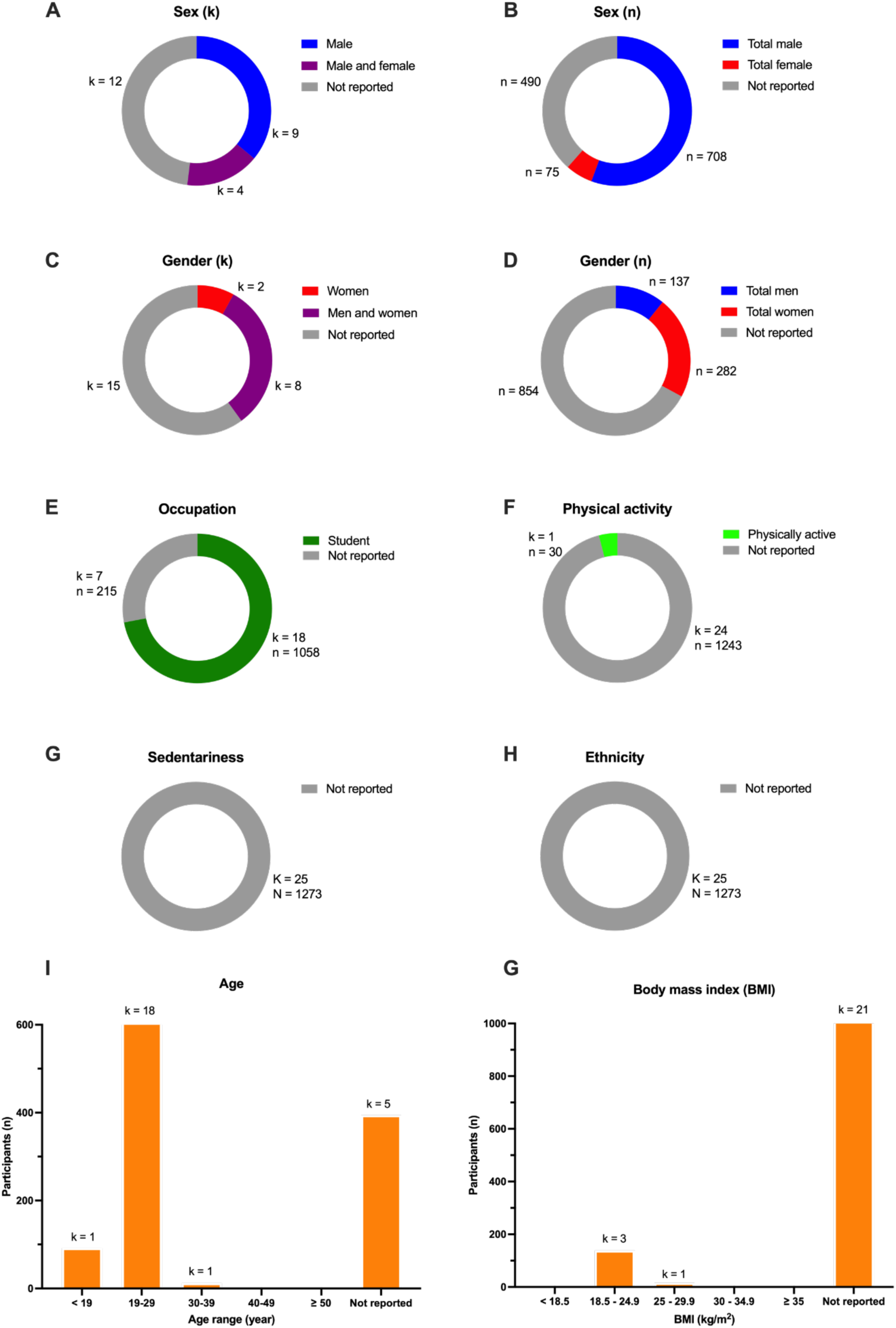
(A) Number of female and male participants (n) in the included studies (k). (B) Number of men and women participants (n) in the included studies (k). (C) Number of studies (k) that included the participants’ occupation (n). (D) Number of studies (k) that included the participants’ ethnicity (n). (E) Number of studies (k) that included the participants’ physical activity level (n). (F) Number of studies (k) that included the participants’ sedentariness (n). (G) Participants age. (H) Participants’ body mass index.

Regarding gender, 8% of the studies (k = 2) exclusively recruited women (Carron, 1969; Godwin & Schmidt, 1971). Furthermore, 32% of studies (k = 8) included both men and women (Anguera et al., 2012; Arnett, DeLuccia, & Gilmartin, 2000; Banihosseini, Abdoli, & Kavyani, 2025; Borragan et al., 2016; Branscheidt et al., 2019; Godoi Filho et al., 2025; Takahashi et al., 2006; Zabihhosseinian et al., 2020). Sixty percent of studies (k = 15) did not provide any information about participants’ gender (Alderman, 1965; Apreutesei & Cressman, 2024; Berger & Smith-Hale, 1991; Cotten et al., 1974; Cotten et al., 1972; Dickinson, Medhurst, & Whittingham, 1979; Dwyer, 1984; Hoskens et al., 2022; Khojasteh Moghani et al., 2021; Masters, Poolton, & Maxwell, 2008; Mierau et al., 2009; Nardon et al., 2024; Nunney, 1963; Siekirk, Lai, & Kendall, 2019; Whitley, 1975) (Figure 6C&D). This lack of reporting is particularly notable in older studies when gender considerations were not as systematically integrated into study design as they are nowadays.

Eighteen studies recruited college/university students, while 7 studies did not report participants’ occupations (Figure 6E). Among all the studies, one reported the level of physical activity (Mierau et al., 2009) as presented in Figure 6F. None of the studies reported any information about participants’ sedentariness (Figure 6G), defined as any waking behavior that requires low energy expenditure (Panahi & Tremblay, 2018), or ethnicity (Figure 6H). The mean reported age of participants was 22.4 years. No study included participants above 31 years old. Ages ranged between 16.5 years old (Godoi Filho et al., 2025) and 31 years old (Takahashi et al., 2006). Seventy-six percent of studies (k = 19) included participants aged 16-29 years old. Four percent of studies (k = 1) included participants aged 31 years (Takahashi et al., 2006). Twenty percent of studies (k = 5) did not report participant age at all (Alderman, 1965; Dickinson, Medhurst, & Whittingham, 1979; Godwin & Schmidt, 1971; Nunney, 1963; Whitley, 1975) as presented in Figure 6I. Only 16% of studies (k = 4) reported participants’ body mass index (Dwyer, 1984; Mierau et al., 2009; Nardon et al., 2024; Zabihhosseinian et al., 2020). Eighty-four percent of articles (k = 21) had no report of participants’ body mass index (Figure 6G).

### 3.6 Fatigue effects on motor learning

Figure 7 shows the effect of fatigue on motor learning, as reported and interpreted by the authors of each study. When there was no effect, fatigue was induced by physical exertion in 24% (k = 6) of the studies (Alderman, 1965; Carron, 1969; Cotten et al., 1974; Cotten et al., 1972; Dickinson, Medhurst, & Whittingham, 1979; Masters, Poolton, & Maxwell, 2008). When there was a negative effect of fatigue on motor learning, physical exertion accounted for 40% (k = 10) (Arnett, DeLuccia, & Gilmartin, 2000; Berger & Smith-Hale, 1991; Branscheidt et al., 2019; Dwyer, 1984; Godwin & Schmidt, 1971; Nunney, 1963; Siekirk, Lai, & Kendall, 2019; Takahashi et al., 2006; Whitley, 1975; Zabihhosseinian et al., 2020), and cognitive exertion accounted for 16% (k = 4) (Anguera et al., 2012; Apreutesei & Cressman, 2024; Godoi Filho et al., 2025; Khojasteh Moghani et al., 2021) of instances, respectively. Eight percent of studies (k = 2) showed a positive effect of fatigue on motor learning, and it was after fatigue induced by cognitive exertion (Banihosseini, Abdoli, & Kavyani, 2025; Borragan et al., 2016), and physical exertion (Banihosseini, Abdoli, & Kavyani, 2025). Banihosseini et al. (2025) employed both mental and physical exertion to induce fatigue and found an increase in motor learning inferred via an acuity task. Twelve percent of the studies (k = 3) investigated motor acquisition and did not include retention or transfer tests to assess motor learning (Hoskens et al., 2022; Mierau et al., 2009; Nardon et al., 2024).

**Figure 7.**
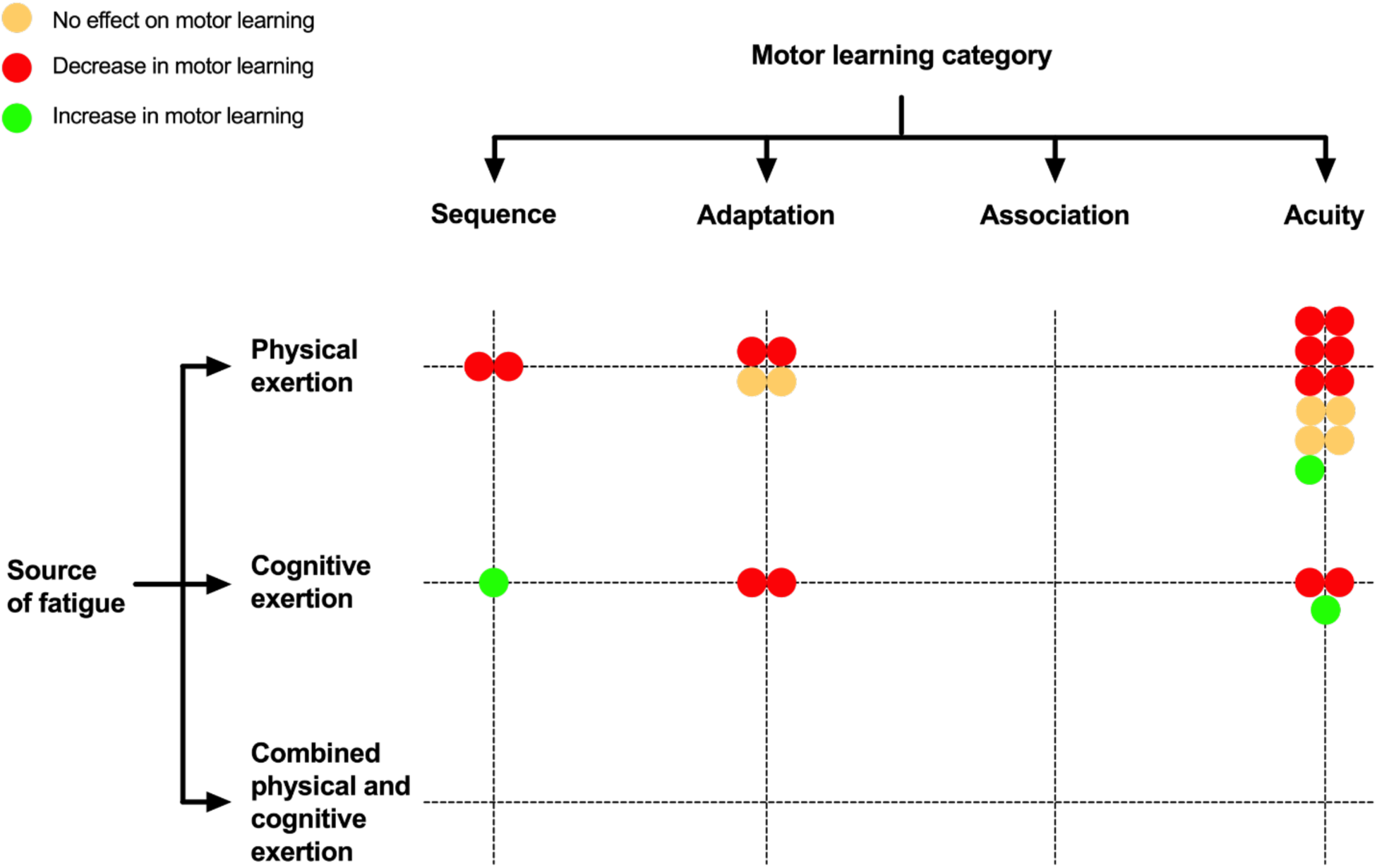
Effect of fatigue on motor learning, based on how the authors of each study interpreted their results. Over the 25 included studies in the scoping review, three did not include retention or transfer tests to assess motor learning and were therefore not represented in this figure. One study included a physical exertion and cognitive exertion protocol to induce fatigue, resulting in the inclusion of two dots (one per protocol) in the figure.

### 3.7 Other sources

An MSc thesis (Datla, 2016) and a book chapter (Taylor, 2013) were included in this scoping review in addition to the experimental studies. The MSc thesis investigated the effects of fatigue on proprioception and motor learning. However, there was no published article associated with the MSc thesis; therefore, no data extraction was conducted on the thesis. The included book chapter (Taylor, 2013) focuses on defining fatigue induced by physical exertion and its effect on neuromuscular function, and then introduces the effect of fatigue induced by physical exertion on neuromuscular function, motor control, and motor performance. The last part of the chapter focuses on the effect of fatigue induced by physical exertion on motor learning, stating that the effect is unclear.

## 4. Discussion

This scoping review aimed to explore the available knowledge on the effects of experimentally induced fatigue on motor learning. This scoping review focused on the populations studied, the methodologies used to induce and measure fatigue, the methodologies used to infer motor learning, as well as the definitions provided for motor learning and fatigue. Twenty-seven sources were systematically identified, including 25 experimental studies, 1 MSc thesis, and 1 book chapter. The main outcomes of this scoping review highlight several critical gaps in the literature: (1) the absence of studies exploring fatigue resulting from a combination of physical and cognitive exertion and the impact on motor learning, coupled with limited research on fatigue induced by cognitive exertion alone, (2) a narrow range of motor learning task categories investigated in laboratory settings, (3) inconsistencies in the definitions of fatigue and motor learning across studies, underscoring the need for standardized operational definitions, (4) a notable absence of studies including middle-aged and older adults, and (5) an inconsistent reporting of participant characteristics, such as sex, gender, ethnicity, physical activity level, sedentariness, and body mass index.

### 4.1 Fatigue

#### 4.1.1 How was fatigue experimentally induced?

This scoping review revealed that the majority of studies investigated fatigue in the physical domain by focusing on fatigue induced by physical exertion. However, the effect of fatigue induced by cognitive exertion remains relatively unclear despite its growing relevance. Fatigue induced by cognitive exertion is increasingly recognized as a distinct construct from fatigue induced by physical exertion (Dong et al., 2022; Russell et al., 2020). Some daily life motor performance experiences, such as typing, require substantially intense and/or prolonged cognitive exertion. This reflects the importance of measuring the impact of fatigue resulting from cognitive exertion on motor learning. However, the current literature on fatigue and motor learning remains limited, leaving a significant gap in studying the effect of fatigue resulting from cognitive exertion on motor learning.

In addition to the limited research on fatigue induced by cognitive exertion, no study investigating motor learning has explored fatigue resulting from a combination of both physical and cognitive exertion. Studying how these combined sources of fatigue influence motor learning is critical, as it reflects real-world scenarios such as firefighting, where individuals must perform motor tasks under high physical and cognitive loads. This area remains unexplored, leaving a significant gap in the literature. Future research should prioritize developing standardized protocols that account for individual differences and explore the mechanisms through which combined physical and cognitive exertion influences motor learning. However, methodological challenges further complicate research in this area (Dong et al., 2022). These challenges underscore the need to introduce innovative methodologies.

Furthermore, only 4 out of the 25 studies reported the time interval between the fatiguing task and the motor learning task. This is a critical piece of information, as the timing of the motor learning task relative to fatigue induction can significantly influence outcomes. While the recovery dynamics following cognitively induced fatigue remain poorly understood, it is well known that muscle fatigue tends to recover rapidly (Doyle-Baker et al., 2018; Pageaux et al., 2015). Therefore, without reporting this time interval, it becomes difficult to interpret the extent to which fatigue was present during motor task practice.

#### 4.1.2 What criteria were used to verify the successful fatigue induction?

Fatigue alters cognitive and physical functioning and can manifest subjectively and objectively (e.g., Mangin & Pageaux, 2025). Additionally, interindividual differences can lead to unique manifestations of fatigue (Dong et al., 2022; Wilson et al., 2021). Therefore, capturing both manifestations is essential for accurately characterizing the nature and extent of fatigue induced by a given task. Given the absence of a universally accepted gold standard for measuring fatigue (Behrens et al., 2023; Dong et al., 2022; Mangin & Pageaux, 2025) it is essential to incorporate both subjective and objective manifestations to comprehensively assess fatigue (Mangin & Pageaux, 2025).

Among all the studies in this scoping review, 12% of the included designs (k = 3) verified the presence of fatigue by measuring both objective (behavioral) and subjective manifestations (Apreutesei & Cressman, 2024; Banihosseini, Abdoli, & Kavyani, 2025; Borragan et al., 2016). Another study (Hoskens et al., 2022) used both subjective and objective (behavioral and physiological) criteria to verify the successful induction of fatigue induced by cognitive exertion; however, this study did not include retention or transfer tests to infer motor learning, and they merely studied motor acquisition. Integrating physiological measures alongside subjective and behavioral indicators can provide a more comprehensive assessment of fatigue and improve the translational relevance of future studies (e.g., Finsterer, 2012; Guffey et al., 2012; Ražanskas et al., 2015). In three studies (Godoi Filho et al., 2025; Khojasteh Moghani et al., 2021; Nunney, 1963), the presence of fatigue was confirmed using self-reported measures of fatigue exclusively, and the rest of the studies only used a behavioral criterion to verify successful fatigue induction. Moreover, none of the studies utilized a self-reported measure of effort. Given the interplay between the perception of effort with motor learning (Ghafari Goushe et al., 2024) and fatigue (Pageaux & Lepers, 2016), measuring the perception of effort can provide valuable insights into the mechanisms through which fatigue influences motor learning.

#### 4.1.3 Towards a unifying definition of fatigue

As highlighted in the previous section, inconsistencies in the definition of fatigue and the framework used are evident across the literature (see supplementary material 4). These inconsistencies are reflected in the heterogeneous methods used and the variability of findings reported. While the literature still does not agree on a definition of fatigue across disciplines (Behrens et al., 2023; Kluger, Krupp, & Enoka, 2013), this scoping review provides a valuable opportunity to clarify and standardize the conceptualization of fatigue.

The use of varied concepts for the same phenomenon, both within and across different research fields, impedes interdisciplinary collaboration and can lead to confusion and miscommunication (Mangin & Pageaux, 2025; Skau, Sundberg, & Kuhn, 2021). Researchers investigating fatigue from a physiological standpoint often overlook its subjective manifestations, whereas those approaching it from a psychological perspective frequently neglect its objective indicators. This holds true even when the authors do not provide an explicit definition of fatigue, but the conceptual framework can nonetheless be inferred from their writing. A comprehensive definition of fatigue should avoid being overly discipline-specific to ensure cross-disciplinary applicability and enhance generalisability (Briand et al., 2025). At the same time, the definition should specify the primary measure of interest (Briand et al., 2025), enabling the monitoring of both the objective and subjective manifestations of fatigue, as well as the origin of fatigue (e.g., depletion of resources).

Mangin and Pageaux (2025) defined fatigue as a psychobiological phenomenon induced by prolonged engagement in physical and/or cognitive exertion or illness, and characterized by depletion or inaccessibility of resources. The presence of fatigue can be assessed through both subjective and objective measures. Subjectively, fatigue manifests as increased feelings of tiredness or exhaustion, often accompanied by an unwillingness or inability to continue the current task or initiate new ones (Boksem & Tops, 2008; Müller & Apps, 2019; Pageaux & Lepers, 2016). Objectively, if the level of engagement (i.e., effort) in a task remains constant, fatigue impairs physical and cognitive performance. The performance declines may or may not be linked to impairments in physiological responses to the task (Behrens et al., 2023; Boksem & Tops, 2008; Gandevia, 2001; Mangin & Pageaux, 2025). When objective manifestations of fatigue are identified, they may reflect alterations at multiple levels of the neuromuscular system, including cortical, spinal, and peripheral components, without necessarily being attributable to a single underlying mechanism. This multilevel nature of fatigue-related alterations is particularly evident when fatigue is induced by tasks involving a substantial physical component. In the presence of fatigue, performance may still be maintained through compensatory mechanisms, such as increasing the effort invested in the task (Wright & Mlynski, 2019). We propose that future research interested in a multidisciplinary investigation of the fatigue phenomenon uses such a definition and description. Using such a definition and description would likely unify the different frameworks of fatigue identified in this scoping review: depletion of resources, decrement in performance and subjective feeling of tiredness.

### 4.2 Motor learning

#### 4.2.1 What task categories were used to study motor learning?

The study of motor learning often relies on a limited set of laboratory-based tasks, which simplify the vast diversity of real-world motor behaviors into controlled, repeatable experimental categories (Krakauer et al., 2019). While these task categories provide valuable insights, a critical question concerns how interchangeable they are when it comes to drawing general conclusions about motor learning. This raises important considerations about the generalizability of results and whether insights from one task can truly apply to the broader spectrum of motor learning (Krakauer et al., 2019).

More than half of the studies included in this scoping review employed an acuity task to infer motor learning. However, none of the articles involved the association task category. Few studies investigated adaptation and sequence task categories. This narrow focus can limit the generalizability of findings. To address this gap, future research should prioritize the investigation of sequence, adaptation, and association task categories. The use of diverse motor tasks is not only essential for enhancing ecological validity but also for improving the replicability of results (Ranganathan et al., 2021). When a result is observed within a single task category, future studies should aim to either directly or conceptually replicate the finding (Derksen & Morawski, 2022). Direct replication serves to confirm the reliability of the effect within the same task category, helping to determine whether the observed effect is valid and/or reliable. In contrast, conceptual replication is crucial for evaluating the generalisability of the effect across different task categories, thereby shedding light on its broader relevance to motor learning beyond the confines of a specific experimental context.

#### 4.2.2 What methods were used to infer motor learning?

Since the performance level achieved during acquisition may not always accurately reflect motor learning, it is critical to assess performance in post-practice phases using retention and/or transfer tests (Christina, 1997; Kantak & Winstein, 2012). A retention test principally measures performance persistence over time. A transfer test mainly measures the generalizability of performance (Christina, 1997; Kantak & Winstein, 2012). Both retention and transfer tests can be used to infer motor learning; however, they serve different purposes depending on the specific aspect of learning being assessed, i.e., persistence over time or generalizability (Christina, 1997; Kantak & Winstein, 2012). According to the traditional definition, motor learning refers to long-lasting improvements in the capability to respond to the demands of a motor task (Schmidt, 1991). This definition inherently requires the inclusion of a delayed retention test to verify that performance improvements are not merely transient but reflect long-lasting changes. Without a retention test, observed gains in performance may simply represent short-term improvements rather than learning (Schmidt, 1991). Our findings indicate that the majority of studies in this scoping review included a retention test, which is a crucial component for assessing motor learning.

Beyond assessing the persistence aspect of motor learning, studies also employ transfer tests to evaluate generalizability of motor learning (Seidler, 2010). Generalization refers to the extent to which a learned skill can be expressed under novel conditions, for example, with a different effector (e.g., transferring from the dominant to the non-dominant hand; Chase & Seidler, 2008), in a new spatial context (e.g., a different workspace; Shadmehr & Mussa-Ivaldi, 1994), or using a different mode of movement or a variation of a task (e.g., transferring from a continuous tracking task to a discrete pointing task; Abeele & Bock, 2003). Transfer tests assess the learner’s capability to generalize what has been learned to performance situations and contexts that were not experienced during practice (Magill & Anderson, 2010; Schmidt, 1991).

#### 4.2.3 Motor learning definition

Of the 25 studies included, only 5 defined motor learning, and among all the definitions, only one (Godoi Filho et al., 2025) captured its core characteristic, i.e., the long-lasting improvement in performance. A unified definition is critical, as it largely determines how we study and assess the variable of interest (e.g., Christina, 1997). A comprehensive definition of motor learning must encompass two key characterizations of the motor learning process (Christina, 1997): (1) it involves a set of processes that lead to long-lasting improvements in an individual’s capacity to meet the task requirements, and (2) it is inferred from long-term performance enhancements that result from practice or experience (Christina, 1997). Schmidt and Lee (1991) defined motor learning as a set of processes associated with practice or experience leading to long-lasting improvement in the capacity to respond to task demands (p. 51). This definition captures the key characteristics of motor learning. Indeed, Schmidt and Lee (1991) used the term “relatively permanent” to refer to the persistent aspect of improved performance, emphasizing the need for a delayed retention test. We encourage future research to use such an accepted definition, while also explicitly reporting the definition.

### 4.3 What populations were investigated?

The findings of this scoping review revealed a significant limitation in the demographic representation of participants across the included studies. All participants fell within the age range of 16.5-31 years, highlighting a critical gap in understanding how fatigue impacts motor learning in middle-aged and older adults. Furthermore, it is important to dissociate middle-aged adults from older adults due to age-related differences in fatigue (e.g., Meng, Hale, & Friedberg, 2010) and motor learning (e.g., Boyd, Vidoni, & Siengsukon, 2008). Investigating how fatigue affects motor learning across the lifespan will provide valuable insights into age-specific challenges and inform targeted interventions for older populations (e.g., Zengarini et al., 2015). Future studies should expand their participant pools to include these unrepresented age groups.

Additionally, the reporting of participant characteristics was inconsistent and often absent. More than half of the studies failed to report any information about participants’ sex and gender. Given the relatively recent findings demonstrating the influence of these factors on motor learning and fatigue (e.g., Albert et al., 2006; Demura, Nakada, & Nagasawa, 2008; Hunter, 2009), we strongly encourage future studies to report such information. Moreover, reporting information about participants’ physical activity level and sedentariness that can influence fatigue (e.g., Bogdanis, 2012; Panahi & Tremblay, 2018) and motor learning (e.g., Holfelder & Schott, 2014; Kuo et al., 2023) was overlooked. Research suggests that individuals who are more physically active exhibit greater hippocampal, prefrontal cortex, and basal ganglia volumes, stronger white matter integrity, and superior executive and memory function (e.g., Erickson, Hillman, & Kramer, 2015). Furthermore, randomized controlled trials offer evidence linking moderate-to-vigorous physical activity to cognitive improvements, including enhanced processing speed, memory, and executive function (Erickson et al., 2019). Given the critical role of cognition in motor learning, accounting for participants’ physical activity levels and sedentariness is relevant and important to report. Furthermore, none of the studies included information about participants’ ethnicity, limiting the generalizability of findings across diverse populations (Cordero et al., 2012; Meng, Hale, & Friedberg, 2010).

Only 16% of studies (k = 4) reported participants’ body mass index (BMI), despite its potential relevance to physical and cognitive performance (Dewi, Rimawati, & Purbodjati, 2021; Kim, Kim, & Park, 2016). The BMI is widely used in population-based studies due to its broad acceptance as a tool for categorizing body mass in relation to health risks (Nuttall, 2015). However, it has become increasingly evident that BMI is a poor indicator of actual body fat percentage (Nuttall, 2015). More importantly, it fails to provide information on fat distribution across different regions of the body (Nuttall, 2015). Therefore, whenever feasible, we recommend including additional anthropometric measures, such as percent lean and fat mass, waist circumference, and similar metrics to obtain a more accurate and comprehensive assessment of body composition.

These gaps underscore the need for more comprehensive and standardized reporting of demographic and anthropometric data in future research. To address these gaps, future studies should prioritize inclusivity by recruiting participants from a wider range of ages, sexes, genders, and ethnic backgrounds. Reporting key demographic and physiological characteristics, such as BMI, physical activity level, and sedentariness, should also become standard practice. By doing so, researchers can ensure that findings are more representative of the broader population and provide a stronger foundation to investigate the effects of fatigue on motor learning across diverse groups.

### 4.4 What conclusions did the included studies draw regarding the effects of experimentally induced fatigue on motor learning?

Although our scoping review did not aim to conduct a meta-analysis due to the methodological heterogeneity by which fatigue and motor learning were studied in the literature, it is still possible to discuss the findings as reported by the original authors (see Figure 7 and Supplementary Table S2). The effects of fatigue on motor learning varied depending on the type of exertion used to induce fatigue. When fatigue was induced by physical exertion, the included studies reported either a negative effect (k = 10), no effect (k = 6), or a positive effect (k = 1). When fatigue was induced by cognitive exertion, the included studies reported either a negative effect (k = 4) or a positive effect (k = 2). When sequence tasks were employed, the included studies reported either a negative effect (k = 2) or a positive effect (k = 1). When adaptation tasks were employed, the included studies reported either a negative effect (k = 4) or no effect (k = 2). When acuity tasks were employed, the included studies reported a negative effect (k = 8), a positive effect (k = 2), and no effect (k = 4).

This finding suggests that the relationship between fatigue and motor learning may be more nuanced than previously thought, with potential moderating factors such as the source of fatigue and the specific motor learning tasks employed influencing outcomes. Mechanistically, fatigue may function as an enhanced source of sensory uncertainty that shifts reliance toward prior beliefs in adaptive motor control and learning (Körding & Wolpert, 2004). Furthermore, fatigue induced by physical and cognitive exertion might differentially impact the fast process responsible for rapid skill acquisition or the slow process crucial for long-term skill retention (Smith, Ghazizadeh, & Shadmehr, 2006). These results underscore the need for future research to explore a broader range of motor learning categories and fatigue-induction methods. By diversifying both the motor learning task categories and fatigue-induction methods used in future studies, researchers can gain a more comprehensive understanding of how fatigue impacts motor learning across different contexts and populations.

### 4.5 Limitations

This review focused on experimentally induced fatigue, given the need to isolate the effects of fatigue on motor learning under both physical and cognitive exertion. While this approach allowed us to examine task-induced fatigue, it excluded studies in which fatigue arises from pathological conditions. Future scoping reviews should extend this work by specifically addressing fatigue associated with pathological states. Additionally, it is important to note that fatigue is highly task-specific, even within the same domain. For example, the underlying mechanisms of fatigue differ between whole-body versus isolated muscle exercise (e.g., Sidhu, Cresswell, & Carroll, 2013) and between different cognitive tasks, such as the n-back task versus the Stroop task (Smith et al., 2019). In our review, we categorized fatigue studies by domain rather than by task. This decision was driven by the limited number of studies available for each specific task. As the literature grows, future reviews should aim to examine task-specific differences in fatigue mechanisms to provide a more nuanced understanding of how fatigue influences motor learning across diverse contexts.

### 4.6 Conclusions and recommendations

This scoping review highlights the current state of research on the effects of fatigue on motor learning, revealing critical gaps in the literature. Inconsistencies in methodology and a lack of clear definitions of fatigue and motor learning are evident in the literature, highlighting the need for standardized, operational definitions to improve conceptual clarity and cross-study comparisons. Furthermore, the overreliance on one motor learning task category and the underrepresentation of other categories may limit the direct and conceptual replication of studies. Finally, the narrow demographic representation, with participants predominantly aged 16.5-31 years, and inconsistent reporting of key participant characteristics, restricts the generalizability of findings to broader populations.

Future studies should adopt a comprehensive definition of fatigue and avoid being overly discipline-specific to ensure cross-disciplinary applicability. At the same time, the definition should specify the primary measure of interest, enabling the monitoring of both the objective and subjective manifestations of fatigue. Standardized protocols for fatigue induction and motor learning assessment should be developed to enhance ecological validity and reproducibility. Moreover, future studies should expand their participant pools to include middle-aged and older adults, as well as individuals from diverse demographic backgrounds. Consistent reporting of key participant characteristics should be prioritized.

