## supplementary material 1 for "Experimentally induced fatigue and motor learning: A scoping review"

### Supplementary Material 1
### *detailed search equations*

**Search #1**

Date search : 2021-11-10
The results: 2709 – 834 (duplicates) = 1887 (without duplicates) + 2 theses (see at the end)
œ

**Search terms**

| Concepts | Fatigue | Motor learning |
|---|---|---|
| Keywords | "mental fatigue" OR "physical fatigue" OR "central fatigue" OR "muscle fatigue" OR "muscular fatigue" OR "cognitive fatigue" OR "neural fatigue" OR "peripheral fatigue" OR "neuromuscular fatigue" OR "mental exertion" OR "physical exertion" OR depletion OR "rating of perceived exertion" | "motor learning" OR "motor skill" OR "motor skills" OR "motor adaptation" OR "motor acuity" OR "motor coordination" OR "de novo learning" OR "motor control" OR "motor sequence" OR "motor behavior" OR "motor behaviour" "sensorimotor learning" OR "motor competence" OR "motor competences" |
| **Databases** | **Index terms** | **Index terms** |
| Medline (Ovid) 1946- | fatigue/ or mental fatigue/<br><br>Muscle Fatigue/ | motor skills/ |
| Embase (Ovid) 1974- | fatigue/ or exhaustion/ or muscle fatigue/ | motor learning/<br><br>motor control/ or motor coordination/ |
| PsycInfo (Ovid) 1806- | fatigue/ | Motor Coordination/ or Motor Skills/ or Motor Control/ or Perceptual Motor Coordination/ or Perceptual Motor Learning/ or Gross Motor Skill Learning/ or Fine Motor Skill Learning/ |

| | | |
|---|---|---|
| SportDiscus with Full Text (Ebsco) 1975- | DE "FATIGUE"<br>DE "MENTAL fatigue" | DE "MOTOR learning" OR DE "PERCEPTUAL motor learning" OR DE "MOTOR ability" OR DE "MOVEMENT education" |
| CINAHL Plus with Full Text (Ebsco) 1937- | (MH "Fatigue") OR (MH "Mental Fatigue") OR (MH "Muscle Fatigue") | (MH "Motor Skills") |
| ERIC (ProQuest) 1966- | MAINSUBJECT.EXACT("Fatigue (Biology)") | MAINSUBJECT.EXACT("Perceptual Motor Learning") OR MAINSUBJECT.EXACT("Movement Education") OR MAINSUBJECT.EXACT("Perceptual Motor Coordination") OR MAINSUBJECT.EXACT("Multisensory Learning") OR MAINSUBJECT.EXACT("Psychomotor Skills") |
| Web of Science (Clarivate) 1945- | - | - |
| Dissertations & Theses Global (ProQuest) | - | - |

Database(s): **Ovid MEDLINE(R) ALL** 1946 to November 09, 2021
Search Strategy:

| # | Searches | Results |
|---|---|---|
| 1 | ("mental fatigue" or "physical fatigue" or "central fatigue" or "muscle fatigue" or "muscular fatigue" or "cognitive fatigue" or "neural fatigue" or "peripheral fatigue" or "neuromuscular fatigue" or "mental exertion" or "physical exertion" or depletion or "rating of perceived exertion").ab,kf,ti. | 167626 |
| 2 | fatigue/ or mental fatigue/ | 32793 |
| 3 | Muscle Fatigue/ | 8756 |
| 4 | 1 or 2 or 3 | 202790 |

| | | |
|---|---|---|
| 5 | ("motor learning" or "motor skill" or "motor skills" or "motor adaptation" or "motor acuity" or "motor coordination" or "de novo learning" or "motor control" or "motor sequence" or "motor behavior" or "motor behaviour sensorimotor learning" or "motor competence" or "motor competences").ab,kf,ti. | 39231 |
| 6 | motor skills/ | 25573 |
| 7 | 5 or 6 | 57548 |
| 8 | 4 and 7 | 823 |
| 9 | limit 8 to humans | 549 |
| 10 | limit 9 to english language | 533 |

Database(s): **Embase** 1974 to 2021 November 09
Search Strategy:

| # | Searches | Results |
|---|---|---|
| 1 | ("mental fatigue" or "physical fatigue" or "central fatigue" or "muscle fatigue" or "muscular fatigue" or "cognitive fatigue" or "neural fatigue" or "peripheral fatigue" or "neuromuscular fatigue" or "mental exertion" or "physical exertion" or depletion or "rating of perceived exertion").ab,kw,ti. | 210354 |
| 2 | fatigue/ or exhaustion/ or muscle fatigue/ | 249050 |
| 3 | 1 or 2 | 448553 |
| 4 | ("motor learning" or "motor skill" or "motor skills" or "motor adaptation" or "motor acuity" or "motor coordination" or "de novo learning" or "motor control" or "motor sequence" or "motor behavior" or "motor behaviour sensorimotor learning" or "motor competence" or "motor competences").ab,kw,ti. | 51837 |
| 5 | motor learning/ | 2646 |
| 6 | motor control/ or motor coordination/ | 29038 |
| 7 | 4 or 5 or 6 | 65281 |
| 8 | 3 and 7 | 1530 |
| 9 | limit 8 to human | 1138 |
| 10 | limit 9 to english language | 1118 |
| 11 | limit 10 to embase | 792 |

Database(s): **APA PsycInfo** 1806 to November Week 1 2021
Search Strategy:

| # | Searches | Results |
|---|---|---|

| | | |
|---|---|---|
| 1 | ("mental fatigue" or "physical fatigue" or "central fatigue" or "muscle fatigue" or "muscular fatigue" or "cognitive fatigue" or "neural fatigue" or "peripheral fatigue" or "neuromuscular fatigue" or "mental exertion" or "physical exertion" or depletion or "rating of perceived exertion").ab,ti,tw. | 11879 |
| 2 | fatigue/ | 10076 |
| 3 | 1 or 2 | 20506 |
| 4 | ("motor learning" or "motor skill" or "motor skills" or "motor adaptation" or "motor acuity" or "motor coordination" or "de novo learning" or "motor control" or "motor sequence" or "motor behavior" or "motor behaviour sensorimotor learning" or "motor competence" or "motor competences").ab,ti,tw. | 27007 |
| 5 | Motor Coordination/ or Motor Skills/ or Motor Control/ or Perceptual Motor Coordination/ or Perceptual Motor Learning/ or Gross Motor Skill Learning/ or Fine Motor Skill Learning/ | 17316 |
| 6 | 4 or 5 | 34987 |
| 7 | 3 and 6 | 331 |
| 8 | limit 7 to human | 203 |
| 9 | limit 8 to english | 203 |

**SPORTDiscus with Full Text - EBSCO**

| # | Question | Résultats |
|---|---|---|
| S7 | S3 AND S6<br>Opérateurs de restriction - Relu par un comité de lecture; Langue: English; Type de publication: Academic Journal<br>Opérateurs d'expansion - Appliquer des sujets équivalents<br>Modes de recherche - Booléen/Phrase | 286 |
| S6 | S4 OR S5 | 28,735 |
| S5 | DE "MOTOR learning" OR DE "PERCEPTUAL motor learning" OR DE "MOTOR ability" OR DE "MOVEMENT education" | 24,179 |
| S4 | TI ( "motor learning" OR "motor skill" OR "motor skills" OR "motor adaptation" OR "motor acuity" OR "motor coordination" OR "de novo learning" OR "motor control" OR "motor sequence" OR "motor behavior" OR "motor behaviour" "sensorimotor learning" OR "motor competence" OR "motor competences" ) OR AB ( "motor learning" OR "motor skill" OR "motor skills" OR "motor adaptation" OR "motor acuity" OR "motor coordination" OR "de novo learning" OR "motor control" OR "motor sequence" OR "motor behavior" OR "motor behaviour" "sensorimotor learning" OR "motor competence" OR "motor competences" ) | 10,462 |
| S3 | S1 OR S2 | 16,954 |

| | | |
|---|---|---|
| S2 | DE "FATIGUE" OR DE "MENTAL fatigue" | 10,774 |
| S1 | TI ( "mental fatigue" OR "physical fatigue" OR "central fatigue" OR "muscle fatigue" OR "muscular fatigue" OR "cognitive fatigue" OR "neural fatigue" OR "peripheral fatigue" OR "neuromuscular fatigue" OR "mental exertion" OR "physical exertion" OR depletion OR "rating of perceived exertion" ) OR AB ( "mental fatigue" OR "physical fatigue" OR "central fatigue" OR "muscle fatigue" OR "muscular fatigue" OR "cognitive fatigue" OR "neural fatigue" OR "peripheral fatigue" OR "neuromuscular fatigue" OR "mental exertion" OR "physical exertion" OR depletion OR "rating of perceived exertion" ) | |

**CINAHL Plus with Full Text - EBSCO**

| # | Question | Résultats |
|---|---|---|
| S7 | S3 AND S6<br>Opérateurs de restriction - Langue anglaise; Relu par un comité de lecture<br>Opérateurs d'expansion - Appliquer des sujets équivalents<br>Modes de recherche - Booléen/Phrase | 214 |
| S6 | S4 OR S5 | 18,100 |
| S5 | (MH "Motor Skills") | 11,726 |
| S4 | TI ( "motor learning" OR "motor skill" OR "motor skills" OR "motor adaptation" OR "motor acuity" OR "motor coordination" OR "de novo learning" OR "motor control" OR "motor sequence" OR "motor behavior" OR "motor behaviour" "sensorimotor learning" OR "motor competence" OR "motor competences" ) OR AB ( "motor learning" OR "motor skill" OR "motor skills" OR "motor adaptation" OR "motor acuity" OR "motor coordination" OR "de novo learning" OR "motor control" OR "motor sequence" OR "motor behavior" OR "motor behaviour" "sensorimotor learning" OR "motor competence" OR "motor competences" ) | 9,741 |
| S3 | S1 OR S2 | 34,177 |
| S2 | (MH "Fatigue") OR (MH "Mental Fatigue") OR (MH "Muscle Fatigue") | 23,394 |
| S1 | TI ( "mental fatigue" OR "physical fatigue" OR "central fatigue" OR "muscle fatigue" OR "muscular fatigue" OR "cognitive fatigue" OR "neural fatigue" OR "peripheral fatigue" OR "neuromuscular fatigue" OR "mental exertion" OR "physical exertion" OR depletion OR "rating of perceived exertion" ) OR AB ( "mental fatigue" OR "physical fatigue" OR "central fatigue" OR "muscle fatigue" OR "muscular fatigue" OR "cognitive fatigue" OR "neural fatigue" OR "peripheral fatigue" OR "neuromuscular fatigue" OR "mental exertion" OR "physical exertion" OR depletion OR "rating of perceived exertion" ) | 13,248 |

**ERIC - Proquest**

| Search | Results |
|---|---|
| ((noft("mental fatigue" OR "physical fatigue" OR "central fatigue" OR "muscle fatigue" OR "muscular fatigue" OR "cognitive fatigue" OR "neural fatigue" OR "peripheral fatigue" OR "neuromuscular fatigue" OR "mental exertion" OR "physical exertion" OR depletion OR "rating of perceived exertion") OR MAINSUBJECT.EXACT("Fatigue (Biology)")) AND PEER(yes)) AND ((noft("motor learning" OR "motor skill" OR "motor skills" OR "motor adaptation" OR "motor acuity" OR "motor coordination" OR "de novo learning" OR "motor control" OR "motor sequence" OR "motor behavior" OR "motor behaviour" "sensorimotor learning" OR "motor competence" OR "motor competences") OR (MAINSUBJECT.EXACT("Perceptual Motor Learning") OR MAINSUBJECT.EXACT("Movement Education") OR MAINSUBJECT.EXACT("Perceptual Motor Coordination") OR MAINSUBJECT.EXACT("Multisensory Learning") OR MAINSUBJECT.EXACT("Psychomotor Skills"))) AND PEER(yes)) | **16** |

**Web of Science Core Collection**

"mental fatigue" OR "physical fatigue" OR "central fatigue" OR "muscle fatigue" OR "muscular fatigue" OR "cognitive fatigue" OR "neural fatigue" OR "peripheral fatigue" OR "neuromuscular fatigue" OR "mental exertion" OR "physical exertion" OR depletion OR "rating of perceived exertion" **(Topic)** and "motor learning" OR "motor skill" OR "motor skills" OR "motor adaptation" OR "motor acuity" OR "motor coordination" OR "de novo learning" OR "motor control" OR "motor sequence" OR "motor behavior" OR "motor behaviour" "sensorimotor learning" OR "motor competence" OR "motor competences" **(Topic)** and English **(Languages)**

Results : 665

-

**Dissertations & Theses Global (ProQuest)**

Search

ti(Fatigue) AND ti(motor learning) > 2010 = 2

1- Metwali, M. (2019). *Motor variablity, task performance, and muscle fatigue during training of a repetitive lifting task: Adapting motor learning topics to occupational ergonomics research* (Order No. 13862704). Available from ProQuest Dissertations & Theses Global. (2306302727). https://www.proquest.com/dissertations-theses/motor-variablity-task-performance-muscle-fatigue/docview/2306302727/se-2?accountid=12543

Low back problems are among the most common nonfatal occupational injuries reported in the United States, and account for substantial healthcare expenditures (e.g., medical care costs) and losses to worker productivity. A strong association has been well-documented between occupational exposure to repetitive trunk motion and low back problems, particularly among workers performing manual material handling (i.e., lifting) activities. A feature of repetitive motion believed important to the development of work-related musculoskeletal disorders (MSDs), including low back problems, is a lack of within-individual, between-cycle variation of physical exposure summary measures, e.g., when observed visually, the cycle-to-cycle motion pattern appears consistent. An active literature has emerged using concepts of motor control to improve ergonomists' understanding of physical exposure variation (i.e., motor variability) arising from individual-level mechanisms during repetitive work.Fundamentally, for any particular individual, the onset of exposure to a repetitive physical activity (i.e., task training) involves a learning process during which motor control strategies are developed to accomplish the task effectively. The cycle-to-cycle variability of motor learning metrics, such as postural and task performance summary measures, has been observed to exponentially decay during task training. From an ergonomics perspective, a temporal reduction in postural variability may lead to greater cumulative loading and physiological fatiguing of the underlying muscle tissues (due to more consistent cycle-to-cycle movements), thus increasing MSD risk over time. However, it is not known if, or to what extent, physical task characteristics (e.g., work pace) modify the temporal behavior of motor variability during training of a repetitive occupational activity. Moreover, the relationships between motor variability, task performance, and muscle fatigue during occupational task training are not well understood.The goal of this dissertation was to present new information concerning occupationally relevant metrics of motor learning during training of a laboratory-simulated, repetitive lifting activity. In this study, participants performed 100 repetitions (i.e., cycles) of the lifting task in each of four experimental sessions (i.e., visits) at different combinations of box load (low or high) and work pace (slow or fast). Three main observations were discussed in this dissertation: (i) participants exhibited a greater temporal reduction in the cycle-to-cycle variability of trunk postural summary measures during training of a heavier-weighted and faster-paced lifting activity (Chapter 3), which may have facilitated increases in the efficiency and repeatability of box movements (Chapter 4), (ii) the cycle-to-cycle variability of the erector spinae (back) muscle activity summary measures increased, but the variability of the multifidus muscle activity summary measures decreased, over time during faster-paced lifting (Chapter 3), and (iii) a greater temporal increase in trunk postural variability (i.e., a more "flexible" trunk movement strategy) was generally associated with lesser electromyographic back muscle fatigue during training of the lifting task (Chapter 5). Collectively, these research findings may open pathways to the development of new task design criteria and ergonomic guidelines to promote motor variability in the workplace and, ultimately, improve workers' musculoskeletal health.

2- Datla, G. (2016). *Effects of local muscle fatigue on proprioception and motor learning* (Order No. 10112524). Available from ProQuest Dissertations & Theses Global. (1794166895). https://www.proquest.com/dissertations-theses/effects-local-muscle-fatigue-on-proprioception/docview/1794166895/se-2?accountid=12543

Background: Muscle fatigue is an exercise induced decline in the ability of muscles to produce force or power. Recent studies showed that decline in proprioception due to fatigue lead to an increasing risk of falls and injury. However, it was unknown whether fatigue-induced proprioception decrease affects skill acquisition and memory consolidation. Purpose: The aim of the study was to investigate the effects of local muscle fatigue on perceptual motor learning in arm positioning task and to compare surface EMG activities in the fatigue and non-fatigue muscle conditions. Two experiments were used to investigate the purpose. In Experiment 1, Methods: 24 healthy young adults (Age: 20-40) were randomly and equally assigned into either control or experiment group. An informed consent was signed prior to the study. Both the groups performed the same task but the experiment group underwent a fatigue protocol (biceps curls with weight of 80% voluntary contraction until fatigue) during the acquisition phase. The task was to place the left forearm on a kinestheiometer and moved the handle to 30, 45, 60 degrees by flexion. All the participants performed 1 block of pre-test, 5 blocks of acquisition phase, 1 block of post-immediate test during the first visit. A delayed retention and bilateral transfer tests were administered 48 hrs after the first visit. Each block had 12 trials. Throughout the task participants were blind folded and were given verbal feedback during the acquisition only. Results: A 2 X 5 (Group vs. Block) ANOVA with repeated measure on Block for acquisition demonstrated both groups decreased total movement error (E) with practice, F (4, 88) = 10.46, p 0.05] or interaction F (1, 22) = 0.00, P> 0.05]. In Experiment 2, Methods: 12 healthy individuals (age 20-40) participated in the experiment that consisted of 6 blocks with 12 trials each. All the participants performed 6 blocks of task and fatigue protocol before every other block. After fatigue protocol participants were made to perform the task immediately without rest but were given 2 minutes rest after each block. Results: One way ANOVA with repeated measure on condition showed a main effect of fatigue for the EMG frequency, $F(1, 22) = 7$, $P<.05$. Where fatigue condition was greater than non-fatigue. The main effect was also detected for the integral EMG (amplitude), $F(1, 12) = 6.14$, $P< .05$, where non fatigue was greater than fatigue. Conclusion: Both the control and experiment group exhibited perceptual motor learning with practice. The fatigue group showed a greater error than the control group in acquisition, retention and transfer. The surface EMG showed increased frequency and decreased integral (amplitude) in the fatigued muscle when compared to non-fatigue condition. In summary, local muscle fatigue had negative effects on perceptual motor acquisition and memory consolidation by degrading proprioception and efficiency on the muscles.

**Search #2**

Research report - update 2025

**Effects of experimentally induced fatigue on motor learning: A scoping review.**

Denis Arvisais

Bibliothécaire
Bibliothèques des sciences de la santé, Université de Montréal

2025-06-10

---

**Updated research**
Here are the results of the 2021 literature search update as of June 10, 2025
1142 – 439 (duplicates) = 703 (without duplicates)

**Previous research**
Search date: 2021-11-10
Results: 2709 – 822 (duplicates) = 1887 (without duplicates) + 2 theses.

-

**Search terms**

| Concepts | Fatigue | Motor learning |
|---|---|---|

| | | |
|---|---|---|
| Keywords | "mental fatigue" OR "physical fatigue" OR "central fatigue" OR "muscle fatigue" OR "muscular fatigue" OR "cognitive fatigue" OR "neural fatigue" OR "peripheral fatigue" OR "neuromuscular fatigue" OR "mental exertion" OR "physical exertion" OR depletion OR "rating of perceived exertion" | "motor learning" OR "motor skill" OR "motor skills" OR "motor adaptation" OR "motor acuity" OR "motor coordination" OR "de novo learning" OR "motor control" OR "motor sequence" OR "motor behavior" OR "motor behaviour" "sensorimotor learning" OR "motor competence" OR "motor competences" |
| **Databases** | **Index terms** | **Index terms** |
| Medline (Ovid) 1946- | fatigue/ or mental fatigue/<br><br>Muscle Fatigue/ | motor skills/ |
| Embase (Ovid) 1974- | fatigue/ or exhaustion/ or muscle fatigue/ | motor learning/<br><br>motor control/ or motor coordination/ |
| PsycInfo (Ovid) 1806- | fatigue/ | Motor Coordination/ or Motor Skills/ or Motor Control/ or Perceptual Motor Coordination/ or Perceptual Motor Learning/ or Gross Motor Skill Learning/ or Fine Motor Skill Learning/ |
| SportDiscus with Full Text (Ebsco) 1930- | DE "FATIGUE"<br>DE "MENTAL fatigue" | DE "MOTOR learning" OR DE "PERCEPTUAL motor learning" OR DE "MOTOR ability" OR DE "MOVEMENT education" |
| CINAHL Complete (Ebsco) 1937- | (MH "Fatigue") OR (MH "Mental Fatigue") OR (MH "Muscle Fatigue") | (MH "Motor Skills") |
| ERIC (ProQuest) 1966- | MAINSUBJECT.EXACT("Fatigue (Biology)") | MAINSUBJECT.EXACT("Perceptual Motor Learning") OR MAINSUBJECT.EXACT("Movement Education") OR MAINSUBJECT.EXACT("Perceptual Motor Coordination") OR MAINSUBJECT.EXACT("Multisensory Learning") OR MAINSUBJECT.EXACT("Psychomotor Skills") |

| | | |
|---|---|---|
| Web of Science (Clarivate)<br>1945- | - | - |
| ProQuest™ Dissertations & Theses Citation Index (Clarivate)<br>1637- | - | - |

#### Search equations

Database(s): **Ovid MEDLINE(R) ALL** 1946 to June 09, 2025
Search Strategy:

| # | Searches | Results |
|---|---|---|
| 1 | ("mental fatigue" or "physical fatigue" or "central fatigue" or "muscle fatigue" or "muscular fatigue" or "cognitive fatigue" or "neural fatigue" or "peripheral fatigue" or "neuromuscular fatigue" or "mental exertion" or "physical exertion" or depletion or "rating of perceived exertion").ab,kf,ti. | 200219 |
| 2 | fatigue/ or mental fatigue/ | 40144 |
| 3 | Muscle Fatigue/ | 9832 |
| 4 | 1 or 2 or 3 | 242344 |
| 5 | ("motor learning" or "motor skill" or "motor skills" or "motor adaptation" or "motor acuity" or "motor coordination" or "de novo learning" or "motor control" or "motor sequence" or "motor behavior" or "motor behaviour sensorimotor learning" or "motor competence" or "motor competences").ab,kf,ti. | 50001 |
| 6 | motor skills/ | 27661 |
| 7 | 5 or 6 | 68968 |
| 8 | 4 and 7 | 985 |
| 9 | exp Animals/ not Humans/ | 5347590 |
| 10 | 8 not 9 | 744 |
| 11 | limit 10 to yr="2021 -Current" | 161 |

Database(s): **Embase** 1974 to 2025 June 09
Search Strategy:

| # | Searches | Results |
|---|---|---|
| 1 | ("mental fatigue" or "physical fatigue" or "central fatigue" or "muscle fatigue" or "muscular fatigue" or "cognitive fatigue" or "neural fatigue" or "peripheral fatigue" or "neuromuscular fatigue" or "mental exertion" or "physical exertion" or depletion or "rating of perceived exertion").ab,kf,ti. | 258669 |
| 2 | fatigue/ or exhaustion/ or muscle fatigue/ | 349920 |
| 3 | 1 or 2 | 593936 |
| 4 | ("motor learning" or "motor skill" or "motor skills" or "motor adaptation" or "motor acuity" or "motor coordination" or "de novo learning" or "motor control" or "motor sequence" or "motor behavior" or "motor behaviour sensorimotor learning" or "motor competence" or "motor competences").ab,kf,ti. | 68203 |

| | | |
|---|---|---|
| 5 | motor learning/ | 5043 |
| 6 | motor control/ or motor coordination/ | 36136 |
| 7 | 4 or 5 or 6 | 83552 |
| 8 | 3 and 7 | 2249 |
| 9 | (exp animal/ or nonhuman/) not exp human/ | 7706827 |
| 10 | 8 not 9 | 1822 |
| 11 | limit 10 to yr="2021 -Current" | 555 |

Database(s): **APA PsycInfo** 1806 to June 2025 Week 1
Search Strategy:

| # | Searches | Results |
|---|---|---|
| 1 | ("mental fatigue" or "physical fatigue" or "central fatigue" or "muscle fatigue" or "muscular fatigue" or "cognitive fatigue" or "neural fatigue" or "peripheral fatigue" or "neuromuscular fatigue" or "mental exertion" or "physical exertion" or depletion or "rating of perceived exertion").ab,id,ti. | 13585 |
| 2 | fatigue/ | 13147 |
| 3 | 1 or 2 | 24745 |
| 4 | ("motor learning" or "motor skill" or "motor skills" or "motor adaptation" or "motor acuity" or "motor coordination" or "de novo learning" or "motor control" or "motor sequence" or "motor behavior" or "motor behaviour sensorimotor learning" or "motor competence" or "motor competences").ab,id,ti. | 30659 |
| 5 | Motor Coordination/ or Motor Skills/ or Motor Control/ or Perceptual Motor Coordination/ or Perceptual Motor Learning/ or Gross Motor Skill Learning/ or Fine Motor Skill Learning/ | 19649 |
| 6 | 4 or 5 | 39186 |
| 7 | 3 and 6 | 387 |
| 8 | exp animals/ | 391565 |
| 9 | 7 not 8 | 267 |
| 10 | limit 9 to yr="2021 -Current" | 60 |

**SPORTDiscus with Full Text - EBSCO**

((XB (( "motor learning" OR "motor skill" OR "motor skills" OR "motor adaptation" OR "motor acuity" OR "motor coordination" OR "de novo learning" OR "motor control" OR "motor sequence" OR "motor behavior" OR "motor behaviour" "sensorimotor learning" OR "motor competence" OR "motor competences" )) OR (DE "MOTOR learning" OR DE "PERCEPTUAL motor learning" OR DE "MOTOR ability" OR DE "MOVEMENT education")) AND (XB (( "mental fatigue" OR "physical fatigue" OR "central fatigue" OR "muscle fatigue" OR "muscular fatigue" OR "cognitive fatigue" OR "neural fatigue" OR "peripheral fatigue" OR "neuromuscular fatigue" OR "mental exertion" OR "physical exertion" OR depletion OR "rating of perceived exertion" )) OR (DE "FATIGUE" OR DE "MENTAL fatigue"))) NOT TI ((rat or rats or mouse or mice or swine or porcine or murine or sheep or lambs or pigs or piglets or rabbit or rabbits or cat or cats or dog or dogs or cattle or bovine or monkey or monkeys or fish* or trout or scallop or scallops or lobster or lobsters or marmoset*))

Relu par un comité de lecture; 31/10/2021 - 09/06/2025
R=62
-

**CINAHL Plus with Full Text - EBSCO**
((XB (( "motor learning" OR "motor skill" OR "motor skills" OR "motor adaptation" OR "motor acuity" OR "motor coordination" OR "de novo learning" OR "motor control" OR "motor sequence" OR "motor behavior" OR "motor behaviour" "sensorimotor learning" OR "motor competence" OR "motor competences" )) OR (MH "Motor Skills")) AND (XB (( "mental fatigue" OR "physical fatigue" OR "central fatigue" OR "muscle fatigue" OR "muscular fatigue" OR "cognitive fatigue" OR "neural fatigue" OR "peripheral fatigue" OR "neuromuscular fatigue" OR "mental exertion" OR "physical exertion" OR depletion OR "rating of perceived exertion" )) OR ((MH "Fatigue") OR (MH "Mental Fatigue") OR (MH "Muscle Fatigue")))) NOT (MH "Animals+" NOT MH "Human")

Relu par un comité de lecture
31/10/2021 - 09/06/2025
R=61
-

**ERIC - Proquest**

((noft("mental fatigue" OR "physical fatigue" OR "central fatigue" OR "muscle fatigue" OR "muscular fatigue" OR "cognitive fatigue" OR "neural fatigue" OR "peripheral fatigue" OR "neuromuscular fatigue" OR "mental exertion" OR "physical exertion" OR depletion OR "rating of perceived exertion") OR MAINSUBJECT.EXACT("Fatigue (Biology)")) AND PEER(yes)) AND ((noft("motor learning" OR "motor skill" OR "motor skills" OR "motor adaptation" OR "motor acuity" OR "motor coordination" OR "de novo learning" OR "motor control" OR "motor sequence" OR "motor behavior" OR "motor behaviour" "sensorimotor learning" OR "motor competence" OR "motor competences") OR (MAINSUBJECT.EXACT("Perceptual Motor Learning") OR MAINSUBJECT.EXACT("Movement Education") OR MAINSUBJECT.EXACT("Perceptual Motor Coordination") OR MAINSUBJECT.EXACT("Multisensory Learning") OR MAINSUBJECT.EXACT("Psychomotor Skills"))) AND PEER(yes))

Limites appliquées
Date entrée:  2021-11-01 - 2025-06-10;
Revu par les pairs
R =3
-

**Web of Science Core Collection**

"mental fatigue" OR "physical fatigue" OR "central fatigue" OR "muscle fatigue" OR "muscular fatigue" OR "cognitive fatigue" OR "neural fatigue" OR "peripheral fatigue" OR "neuromuscular fatigue" OR "mental exertion" OR "physical exertion" OR depletion OR "rating of perceived exertion" **(Topic)** **and** "motor learning" OR "motor skill" OR "motor skills" OR "motor adaptation" OR "motor acuity" OR "motor coordination" OR "de novo learning" OR "motor control" OR "motor sequence" OR "motor behavior" OR "motor behaviour" "sensorimotor learning" OR "motor competence" OR "motor competences" **(All Fields)** **and** 2025 **or** 2024 **or** 2023 **or** 2022 **or** 2021 **(Publication Years)**
NOT
rat or rats or mouse or mice or swine or porcine or murine or sheep or lambs or pigs or piglets or rabbit or rabbits or cat or cats or dog or dogs or cattle or bovine or monkey or monkeys or fish* or trout or scallop or scallops or lobster or lobsters or marmoset* (Title)

R =240

-

**ProQuest™ Dissertations & Theses Citation Index** (Data updated 2025-06-08)

"motor learning" (Title) **and** fatigue (Title)
2021/11/01 to 2025/06/10
R = 0

-

**Search #3**

Research report - Revision

**Effects of Fatigue on motor learning with aging: A scoping review**

Denis Arvisais

Bibliothécaire
Bibliothèques des sciences de la santé, Université de Montréal

2025-11-12

---

**Revised search**

Search date: 2025-11-12

**Deux articles ajoutés à réviser :**

Marr, C. (2023). Service evaluation reviewing the application of the Bayley Scale of Infant and Toddler development in children with severe, complex and atypical Osteogenesis Imperfecta [Conference Abstract]. *JBMR Plus*, *7(Supplement 1)*, 32. https://dx.doi.org/10.1002/jbm4.10718

Osteogenesis Imperfecta is a rare hereditary condition affecting approximately 1 in 20,000 births. Developmental delay is seen in children with the condition. Assessment is key in ensuring appropriate interventions. Although not a disease specific measure, the Bayley Scale of Infant and Toddler Development (BSITD) is used in clinical practice by the Sheffield Children's Hospital team to assess development in children with with severe, complex and atypical Osteogenesis Imperfecta (SCAOI). OBJECTIVE(S): To retrospectively review BSITD assessments, completed in children with SCAOI, who received care at Sheffield Children's NHS Foundation Trust from January 2017 to August 2020, in order to determine if there are any patterns that emerge across the assessments. METHOD(S): A retrospective analysis of BSITD assessments, between January 2017 and August 2020, in children with SCAOI. RESULT(S): A total of 19 BSITD assessments, from 10 patients, were included. 14 of the assessments were with children with type III OI; 5 had type V OI. Mean age was 19 months 17 days. Of the 19 assessments 4 were incomplete, reasons included; fatigue, lack of interest and time limitations. Application of the regression rule was noted in all subsections. Manipulating blocks, squeezing objects, removing lids, instigating play, using words appropriately to make needs known, establishing head control, prone lying and sitting were tasks across the assessments that children repeatedly failed. Only 4 assessment scored above the 50th percentile in the cognitive subsection, with 10 scoring below the 24th percentile. In the language subsection 2 assessments scored above the 50th percentile, with 9 below the 24th percentile. In the motor subsection none of the assessment scores were above the 50th percentile, with 15 assessments below the 24th percentile. CONCLUSION(S): In clinical practice parents of children with SCAOI are told to expect a delay in their child's gross motor acquisition. The service evaluation supports this statement whilst suggesting that within this patient group developmental delay can be observed in all subsections. Further investigation with a larger cohort is warranted to determine, if the BSITD is the appropriate tool to use and establish if the delay observed is a true reflection of children with SCAOI.

Mierau, A., Schneider, S., Abel, T., Askew, C., Werner, S., & Strüder, H. K. (2009). Improved sensorimotor adaptation after exhaustive exercise is accompanied by altered brain activity [Article]. *Physiology and Behavior*, *96*(1), 115–121. https://dx.doi.org/10.1016/j.physbeh.2008.09.002

Acute exercise has been shown to exhibit different effects on human sensorimotor behavior; however, the causes and mechanisms of the responses are often not clear. The primary aim of the present study was to determine the effects of incremental running until exhaustion on sensorimotor performance and adaptation in a tracking task. Subjects were randomly assigned to a running group (RG), a tracking group (TG), or a running followed by tracking group (RTG), with 10 Subjects assigned to each group. Treadmill running velocity was initially set at 2.0 m s(-1) , increasing by 0.5 m s(-1) every 5 min until exhaustion. Tracking consisted of 35 episodes (each 40 s) where the subjects' task was to track a visual target on a computer screen while the Visual feedback was veridical (performance) or left-right reversed (adaptation). Resting electroencephalographic (EEG) activity was recorded before and after each experimental condition (running, tracking, rest). Tracking performance and the final amount of adaptation did not differ between groups. However, task adaptation was significantly faster in RTG compared to TG. In addition, increased alpha and beta power were observed following tracking in TG but not RTG although exhaustive running failed to induce significant changes in these frequency bands. Our results suggest that exhaustive running can facilitate adaptation processes in a manual tracking task. Attenuated cortical activation following tracking in the exercise condition was interpreted to indicate cortical efficiency and exercise-induced facilitation of selective central processes during actual task demands. (c) 2008 Elsevier Inc. All rights reserved.

-

**Updated research**

Here are the results of the 2021 literature search update as of June 10, 2025

1142 – 439 (duplicates) = 703 (without duplicates)

-

**Previous research**

Search date: 2021-11-10

Results: 2709 – 822 (duplicates) = 1887 (without duplicates) + 2 theses.

#### Search terms

| Concepts | Fatigue | Motor learning |
|---|---|---|
| Keywords | "mental fatigue" OR "physical fatigue" OR "central fatigue" OR "muscle fatigue" OR "muscular fatigue" OR "cognitive fatigue" OR "neural fatigue" OR "peripheral fatigue" OR "neuromuscular fatigue" OR "mental exertion" OR "physical exertion" OR depletion OR "rating of perceived exertion" | "motor learning" OR "motor skill" OR "motor skills" OR "motor adaptation" OR "motor acuity" OR "motor coordination" OR "de novo learning" OR "motor control" OR "motor sequence" OR "motor behavior" OR "motor behaviour" "sensorimotor learning" OR "motor competence" OR "motor competences" OR "motor acquisition" |
| **Databases** | **Index terms** | **Index terms** |
| Medline (Ovid) 1946- | fatigue/ or mental fatigue/<br><br>Muscle Fatigue/ | motor skills/ |
| Embase (Ovid) 1974- | fatigue/ or exhaustion/ or muscle fatigue/ | motor learning/<br><br>motor control/ or motor coordination/ |
| PsycInfo (Ovid) 1806- | fatigue/ | Motor Coordination/ or Motor Skills/ or Motor Control/ or Perceptual Motor Coordination/ or Perceptual Motor Learning/ or Gross Motor Skill Learning/ or Fine Motor Skill Learning/ |
| SportDiscus with Full Text (Ebsco) 1930- | DE "FATIGUE"<br>DE "MENTAL fatigue" | DE "MOTOR learning" OR DE "PERCEPTUAL motor learning" OR DE "MOTOR ability" OR DE "MOVEMENT education" |
| CINAHL Complete (Ebsco) 1937- | (MH "Fatigue") OR (MH "Mental Fatigue") OR (MH "Muscle Fatigue") | (MH "Motor Skills") |

| | | |
|---|---|---|
| ERIC (ProQuest) 1966- | MAINSUBJECT.EXACT("Fatigue (Biology)") | MAINSUBJECT.EXACT("Perceptual Motor Learning") OR MAINSUBJECT.EXACT("Movement Education") OR MAINSUBJECT.EXACT("Perceptual Motor Coordination") OR MAINSUBJECT.EXACT("Multisensory Learning") OR MAINSUBJECT.EXACT("Psychomotor Skills") |
| Web of Science (Clarivate) 1945- | - | - |
| ProQuest™ Dissertations & Theses Citation Index (Clarivate) 1637- | - | - |

**Revised Medline**

Database(s): **Ovid MEDLINE(R) ALL** 1946 to November 11, 2025
Search Strategy:

| # | Searches | Results |
|---|---|---|
| 1 | ("mental fatigue" or "physical fatigue" or "central fatigue" or "muscle fatigue" or "muscular fatigue" or "cognitive fatigue" or "neural fatigue" or "peripheral fatigue" or "neuromuscular fatigue" or "mental exertion" or "physical exertion" or depletion or "rating of perceived exertion").ab,kf,ti. | 205016 |
| 2 | fatigue/ or mental fatigue/ | 40851 |
| 3 | Muscle Fatigue/ | 9979 |
| 4 | 1 or 2 or 3 | 247810 |
| 5 | ("motor learning" or "motor skill" or "motor skills" or "motor adaptation" or "motor acuity" or "motor coordination" or "de novo learning" or "motor control" or "motor sequence" or "motor behavior" or "motor behaviour sensorimotor learning" or "motor competence" or "motor competences" or "motor acquisition").ab,kf,ti. | 51654 |
| 6 | motor skills/ | 27968 |
| 7 | 5 or 6 | 70738 |
| 8 | 4 and 7 | 1016 |
| 9 | exp Animals/ not Humans/ | 5393143 |
| 10 | 8 not 9 | 772 |

##### Revised Embase

Database(s): **Embase** 1974 to 2025 November 11
Search Strategy:

| # | Searches | Results |
|---|---|---|
| 1 | ("mental fatigue" or "physical fatigue" or "central fatigue" or "muscle fatigue" or "muscular fatigue" or "cognitive fatigue" or "neural fatigue" or "peripheral fatigue" or "neuromuscular fatigue" or "mental exertion" or "physical exertion" or depletion or "rating of perceived exertion").ab,kf,ti. | 266488 |
| 2 | fatigue/ or exhaustion/ or muscle fatigue/ | 363710 |
| 3 | 1 or 2 | 614912 |
| 4 | ("motor learning" or "motor skill" or "motor skills" or "motor adaptation" or "motor acuity" or "motor coordination" or "de novo learning" or "motor control" or "motor sequence" or "motor behavior" or "motor behaviour sensorimotor learning" or "motor competence" or "motor competences" or "motor acquisition").ab,kf,ti. | 71009 |
| 5 | motor learning/ | 5470 |
| 6 | motor control/ or motor coordination/ | 37505 |
| 7 | 4 or 5 or 6 | 86618 |
| 8 | 3 and 7 | 2393 |
| 9 | (exp animal/ or nonhuman/) not exp human/ | 7844748 |
| 10 | 8 not 9 | 1954 |

##### Revised APA PsycInfo

Database(s): **APA PsycInfo** 1806 to November 2025 Week 1
Search Strategy:

| # | Searches | Results |
|---|---|---|
| 1 | ("mental fatigue" or "physical fatigue" or "central fatigue" or "muscle fatigue" or "muscular fatigue" or "cognitive fatigue" or "neural fatigue" or "peripheral fatigue" or "neuromuscular fatigue" or "mental exertion" or "physical exertion" or depletion or "rating of perceived exertion").ab,id,ti. | 13789 |

| | | |
|---|---|---|
| 2 | fatigue/ | 13539 |
| 3 | 1 or 2 | 25279 |
| 4 | ("motor learning" or "motor skill" or "motor skills" or "motor adaptation" or "motor acuity" or "motor coordination" or "de novo learning" or "motor control" or "motor sequence" or "motor behavior" or "motor behaviour sensorimotor learning" or "motor competence" or "motor competences" or "motor acquisition").ab,id,ti. | 31177 |
| 5 | Motor Coordination/ or Motor Skills/ or Motor Control/ or Perceptual Motor Coordination/ or Perceptual Motor Learning/ or Gross Motor Skill Learning/ or Fine Motor Skill Learning/ | 19892 |
| 6 | 4 or 5 | 39754 |
| 7 | 3 and 6 | 392 |
| 8 | exp animals/ | 394320 |
| 9 | 7 not 8 | 272 |

**Revised SPORTDiscus with Full Text - EBSCO**((XB (( "motor learning" OR "motor skill" OR "motor skills" OR "motor adaptation" OR "motor acuity" OR "motor coordination" OR "de novo learning" OR "motor control" OR "motor sequence" OR "motor behavior" OR "motor behaviour" "sensorimotor learning" OR "motor competence" OR "motor competences" OR "motor acquisition")) OR (DE "MOTOR learning" OR DE "PERCEPTUAL motor learning" OR DE "MOTOR ability" OR DE "MOVEMENT education")) AND (XB (( "mental fatigue" OR "physical fatigue" OR "central fatigue" OR "muscle fatigue" OR "muscular fatigue" OR "cognitive fatigue" OR "neural fatigue" OR "peripheral fatigue" OR "neuromuscular fatigue" OR "mental exertion" OR "physical exertion" OR depletion OR "rating of perceived exertion" )) OR (DE "FATIGUE" OR DE "MENTAL fatigue"))) NOT TI ((rat or rats or mouse or mice or swine or porcine or murine or sheep or lambs or pigs or piglets or rabbit or rabbits or cat or cats or dog or dogs or cattle or bovine or monkey or monkeys or fish* or trout or scallop or scallops or lobster or lobsters or marmoset*))

**Filtres**

Relu par un comité de lecture

R= 324

**Revised Cinahl Complete**

((XB (( "motor learning" OR "motor skill" OR "motor skills" OR "motor adaptation" OR "motor acuity" OR "motor coordination" OR "de novo learning" OR "motor control" OR "motor sequence" OR "motor behavior" OR "motor behaviour" "sensorimotor learning" OR "motor competence" OR "motor competences" OR "motor acquisition")) OR (MH "Motor Skills")) AND (XB (( "mental fatigue" OR "physical fatigue" OR "central fatigue" OR "muscle fatigue" OR "muscular fatigue" OR "cognitive fatigue" OR "neural fatigue" OR "peripheral fatigue" OR "neuromuscular fatigue" OR "mental exertion" OR "physical exertion" OR depletion OR "rating of perceived exertion" )) OR ((MH "Fatigue") OR (MH "Mental Fatigue") OR (MH "Muscle Fatigue")))) NOT (MH "Animals+" NOT MH "Human")

**Filtres**

Relu par un comité de lecture

R=284

**Revised Eric**

((noft("mental fatigue" OR "physical fatigue" OR "central fatigue" OR "muscle fatigue" OR "muscular fatigue" OR "cognitive fatigue" OR "neural fatigue" OR "peripheral fatigue" OR "neuromuscular fatigue" OR "mental exertion" OR "physical exertion" OR depletion OR "rating of perceived exertion") OR MAINSUBJECT.EXACT("Fatigue (Biology)")) AND PEER(yes)) AND ((noft("motor learning" OR "motor skill" OR "motor skills" OR "motor adaptation" OR "motor acuity" OR "motor coordination" OR "de novo learning" OR "motor control" OR "motor sequence" OR "motor behavior" OR "motor behaviour" "sensorimotor learning" OR "motor competence" OR "motor competences" OR "motor acquisition") OR (MAINSUBJECT.EXACT("Perceptual Motor Learning") OR MAINSUBJECT.EXACT("Movement Education") OR MAINSUBJECT.EXACT("Perceptual Motor Coordination") OR MAINSUBJECT.EXACT("Multisensory Learning") OR MAINSUBJECT.EXACT("Psychomotor Skills"))) AND PEER(yes))

Limité par : Revu par les pairs
R=23

**Revised Web of Science Core Collection**

(TS=("mental fatigue" OR "physical fatigue" OR "central fatigue" OR "muscle fatigue" OR "muscular fatigue" OR "cognitive fatigue" OR "neural fatigue" OR "peripheral fatigue" OR "neuromuscular fatigue" OR "mental exertion" OR "physical exertion" OR depletion OR "rating of perceived exertion" ) AND ALL=("motor learning" OR "motor skill" OR "motor skills" OR "motor adaptation" OR "motor acuity" OR "motor coordination" OR "de novo learning" OR "motor control" OR "motor sequence" OR "motor behavior" OR "motor behaviour" "sensorimotor learning" OR "motor competence" OR "motor competences" OR "motor acquisition" ))

NOT
rat or rats or mouse or mice or swine or porcine or murine or sheep or lambs or pigs or piglets or rabbit or rabbits or cat or cats or dog or dogs or cattle or bovine or monkey or monkeys or fish* or trout or scallop or scallops or lobster or lobsters or marmoset* (Title)

R= 969

-
