## supplementary Material 3 for "Experimentally induced fatigue and motor learning: A scoping review"

### *Table S1*

#### Reasons for exclusion of articles based on the predefined inclusion criteria

| Study | Exclusion Reason | Description |
|---|---|---|
| **Arnett 1999** | Full-text article not found | The full-text article is not found. Only the abstract is available. |
| **Benson 1968** | Wrong intervention | The independent variable was exercise intensity, not fatigue. The exercise intensity was adjusted to each subject based on the heart rate response. There was no validation of successfully inducing fatigue. |
| **Brown 2013** | Wrong outcomes | The aim of this study was to investigate pistol shooting performance in police officers under similar conditions of physical fatigue. There is no motor practice/learning in this study. |
| **Cochrane 1975** | Wrong intervention | This study investigated exercise intensity, not fatigue. There was no validation of successfully inducing fatigue. |
| **Huysmans 2008** | Wrong intervention | This study investigated tracking performance, not learning. |
| **Maruyama 2016** | Article is written in Japanese | The article is in Japanese language. Only the abstract is in English. |
| **Mortimer 2024** | Wrong outcomes | While relevant to motor performance, this article did not include motor learning. |
| **Okubo 1973** | Wrong intervention | This article investigated the effect of learning on fatigue in a driving task. The authors found that motor learning coincided well with lessening of the subjective feeling of fatigue. |
| **Pack 1974** | Wrong intervention | This study involves exercise intensity, not fatigue. There was no validation of successfully inducing fatigue. |
| **Phillips 1968** | Full-text article not found | The full-text article is not found. |
| **Schmidt 1969** | Wrong intervention | This study investigated exercise intensity, not fatigue. There was no validation of successfully inducing fatigue. |
| **Stockard 1974** | Full-text article not found | The full-text article is not found. Only the abstract is available. |
| **Takemi 2023** | Wrong intervention | This article examined fatigue induced concurrently with motor practice, whereas our scoping review focused on studies investigating the effects of fatigue *prior to* subsequent motor practice. |
| **Thomas 1975** | Wrong intervention | This study investigated exercise intensity, not fatigue. There was no validation of successfully inducing fatigue. |

| | | |
|---|---|---|
| **Williams 1975** | Wrong intervention | This study investigated exercise intensity, not fatigue. There was no validation of successfully inducing fatigue. |
| **Williams 1976** | Wrong intervention | This study investigated exercise intensity, not fatigue. There was no validation of successfully inducing fatigue. |
| **Williams 1967** | Wrong intervention | This study investigated exercise intensity, not fatigue. There was no validation of successfully inducing fatigue. |
| **Williams 1979** | Wrong intervention | This study investigated exercise intensity, not fatigue. There was no validation of successfully inducing fatigue. |
