## supplementary material 4 for "Experimentally induced fatigue and motor learning: A scoping review"

### *Table S2*

#### Detailed characteristics of all experimental studies

| Study | Population (n; age ± sd years) | Fatigue induction | | | Motor learning assessment | | | Effect of fatigue on motor learning |
|---|---|---|---|---|---|---|---|---|
| | | Source of fatigue | Fatiguing task | Type of manipulation check | Motor Learning Category | Motor task | Retention | |
| **Alderman (1965)** | Students (120; NR ± NR) | Physical exertion | Continuous circular arm movement (10 min at 120 rpm of 3.45kg resistance) | Objective (Declined performance) | Acuity | Rho learning test Pursuit rotor task | Yes | ↔ |
| **Anguera et al., (2012)** | Students (23; 2.0 ± 1.1) | Cognitive exertion | Spatial working memory fatigue task (20 min) | Objective (Declined performance) | Adaptation | Visuomotor adaptation task | Yes | ↘ |
| **Arnett et al., (2000)** | Students (44; 21.8 ± 2) | Physical exertion | Wingate Test (cycling for 1 min at 70% of the measured mean power output) | Objective (Declined performance) | Acuity | Bachman ladder | NR | ↘ |
| **Berger and Smith-hale, (1991)** | Students (32; 20.1 ± 1.9) | Physical exertion | Leg press (lifting loads of 60% and 80% of maximal strength to exhaustion) | Objective (Declined performance) | Sequence | Jumping task | Yes | ↘ |
| **Borragan et al., (2016)** | NR (23; 23.0 ± 4.1) | Cognitive exertion | Time load dual-back task (16 min) | Objective and subjective (Declined performance and self-report) | Sequence | Serial reaction time task | Yes | ↗ |
| **Branscheidt et al., (2019)** | NR (38; 22.2 ± 2.0) | Physical exertion | Isometric pinch task (sustaining MVC until force drops) | Objective (Declined performance) | Sequence | Isometric pinch task Key pressing task | Yes | ↘ |
| **Carron (1969)** | Students (75; 20.9 ± 2.8) | Physical exertion | Arm ergometer (5 min cycling with a 200kg/min resistance or until exhaustion) | Objective (until exhaustion) | Acuity | Pursuit rotor task | Yes | ↔ |
| **Cotton et al., (1974)** | Students (75; 21.6 ± NR) | Physical exertion | Stool stepping task Reverse curling (30 ascents per min for 7 min by a 23 pound reverse curling) | Objective (until exhaustion) | Adaptation | Mirror target toss test | Yes | ↔ |
| **Cotton et al., (1972)** | Students (75; 21.4 ± NR) | Physical exertion | Stool stepping task (30 ascents per min for 7 min by a 23 pound reverse curling) | Objective (until exhaustion) | Adaptation | Mirror target toss test | Yes | ↔ |
| **Dickinson et al., (1979)** | Students (10; NR ± NR) | Physical exertion | Bicycle ergometer (7 min cycling at 40 rpm) | Objective (until exhaustion) | Acuity | Fitts' reciprocal tapping task | Yes | ↔ |
| **Dwyer (1984)** | Students (80; 22.3 ± 2.7) | Physical exertion | Step up task (stepping 60 times per minute) | Objective (until exhaustion) | Acuity | Bachman ladder | Yes | ↘ |
| **Godoi Filho et al., (2025)** | Students (92; 21.4 ± 1.4) | Cognitive exertion | Stroop task (30 min) | Subjective | Acuity | Visuomotor tracking task | Yes | ↘ |
| **Godwin and Schmidt, (1971)** | Students (64; 16.5 ± 1.74) | Physical exertion | Arm ergometer (cycling with 70.2kg/min at 60 rpm) | Objective (until exhaustion) | Acuity | Sigma task | Yes | ↘ |
| **Khojasteh Moghani et al., (2021)** | Students (44; 21.4 ± 1.4) | Cognitive exertion | Stroop task (60 min) | Subjective (self-report) | Acuity | Force production task | Yes | ↘ |
| **Masters et al., (2008)** | NR (41; 20.5 ± 1.2) | Physical exertion | VO2 max running test | Objective (until exhaustion) | Acuity | Rugby passing | Yes | ↔ |
| **Mierau et al., (2009)** | Students (30; 26 ± 4) | Physical exertion | Incremental running | Objective (until exhaustion) | Adaptation | Manual tracking task | NR | ↗ |
| **Nardon et al., (2024)** | NR (28; 22.4 ± 2.9) | Physical exertion | Isometric contraction | Objective (Declined performance) | Adaptation | Force field adaptation reaching task | NR | ↘ |
| **Nunney (1963)** | Students (80; NR ± NR) | Physical exertion | Bicycle ergometer (5 min at different loads and speeds) | Subjective | Acuity | Snoddy stabilometer Pursuit rotor task | Yes | ↘ |
| **Siekirk et al., (2018)** | Students (22; 24.5 ± NR) | Physical exertion | Standing elbow flexion (elbow flexions at 75-85% of the measured maximal value) | Objective (until exhaustion) | Acuity | Joint position matching task | Yes | ↘ |
| **Takahashi et al., (2006)** | NR (12; 31.0 ± NR) | Physical exertion | Reaching with elastic band (starting at 65% and finishing at 55% maximal strength) | Objective | Adaptation | Force skill adaptation reaching task | Yes | ↔ |
| **Whitley (1975)** | Students (120; NR ± NR) | Physical exertion | Heavy mass (gradual increase) | Objective | Acuity | Foot rotational tracking task | Yes | ↘ |
| **Zabihhosseinian et al., (2020)** | NR (16; 21.1 ± 1.5) | Physical exertion | Neck fatigue (2kg head holding) | Objective | Sequence | Tracing task | Yes | ↘ |
| **Banihosseini et al., (2025)** | Students (32; 29.0 ± 5.6) | Cognitive and physical exertion | Stroop tasks & isometric contraction (30 min & 50% MVC for 2 min) | Objective and subjective (Declined performance and self-report) | Acuity | Tennis ball throwing | Yes | ↗ |
| **Hoskens et al., (2022)** | NR (57; 24.0 ± 5.8) | Cognitive exertion | Victoria Stroop Task | Objective and subjective (Declined performance, EEG, and self-report) | Acuity | Shuffleboard Task | NR | ↘ |
| **Apreutesei and Cressman (2024)** | Students (40; 19.8 ± 1.7) | Cognitive exertion | Time load dual-back task (32 min) | Objective and subjective (Declined performance and self-report) | Adaptation | Visuomotor rotation task | Yes | ↘ |

bpm = beats per minute; MVC = maximal voluntary contraction; kg = kilograms; min = minutes; h = hours; d = days; NR = not reported; ↗ = significant enhancement; ↘ = significant decrease; ↔ = no change.
