## supplementary material 5 for "Experimentally induced fatigue and motor learning: A scoping review"

### *Table S3*

#### Fatigue framework(s) used in each study

Table 2 in the article presents the definitions of fatigue and motor learning found in the literature. Many articles do not explicitly define the term "fatigue". However, it is still possible to infer the conceptual framework adopted by the authors based on their descriptions. In the following table, the first column lists the authors' names and the year of publication. The second column provides the definition of fatigue, if available. The third column contains the authors' descriptions related to fatigue, and in bold the definition within its context. The fourth column presents the inferred framework based on the information provided in the third column.

| STUDY | FATIGUE DEFINITION | SENTENCES IN THE ARTICLES | FRAMEWORK |
|---|---|---|---|
| ***STUDIES INVESTIGATING THE EFFECT OF FATIGUE INDUCED BY PHYSICAL EXERTION*** | | | |
| **ALDERMAN 1965** | - | "Thus the present study is concerned with establishing whether, in performing motor learning tasks, impairment in performance caused by fatigue does or does not cause impairment in motor learning." (p. 131) | Fatigue is conceptualized as an *objective performance decrement* |
| **ARNETT ET AL. 2000** | - | - | |
| **BANIHOSSEINI ET AL. 2025** | - | "The previous studies showed pervasive effect of physiological and mental fatigue on working memory, cognitive and motor performance that requires attention (Schmidt et al., 2018; Lam, 2008; Schmidt, 1975). Fatigue manifests as physical fatigue from prolonged tasks or mental fatigue stemming from extended cognitive engagement (Masters et al., 2008)." (p. 2)<br><br>"Mental fatigue has been shown to impair selective attention, influencing tactical performance in sports (Faber et al., 2012; Badin et al., 2016). Moreover, cognitive fatigue has been associated with enhanced procedural sequence learning by imposing stress on working memory, emphasizing the | Fatigue is conceptualized as a *combined objective performance decrement* and *cognitive-resource depletion*. |

| | | | |
|---|---|---|---|
| | | potential of implicit memory resources (Borragán et al., 2016; Yang et al., 2020).” (p. 2) | |
| **BERGER AND SMITH-HALE 1991** | Inability to generate the required or expected force (Edwards 1981) or any reduction in maximum force-generating capacity (Bigland-Ritchie et al., 1983) (p. 156) | “None of these studies, which examined the effects of fatigue on learning of a gross motor tasks, quantified neuromuscular fatigue in terms of “**an ability to generate the required or expected force” (10) or as “any reduction in maximum force generating capacity”** (3). In fact, fatigue was not induced to a level that was similarly intense for all subjects, and when it was in two studies (6, 7), heart rate of 180 beats per minute was the basis for standardization.” (p. 156) | Fatigue is conceptualized as an *objective motor performance decrement* |
| **BRANSCHEIDT ET AL. 2019** | The degradation of maximal force output induced through voluntary physical exertion of task-relevant muscles (no reference) (p. 1) | “Studies investigating fatigue have made a distinction between fatigue as a cognitive phenomenon and fatigue as a neuromuscular phenomenon (Janet, 2012), although this separation can be blurred at times (Kuppuswamy, 2017). In neurological conditions, for instance, fatigue has been described as an overall state linked to changes in motor cortex excitability (Kuppuswamy et al., 2015). **Here, we use the term fatigue to describe the degradation of maximal force output induced through voluntary physical exertion of task-relevant muscles**.” (p. 1) | Fatigue is conceptualized as an *objective motor performance decrement* |
| **CARRON 1969** | - | “The major research emphasis to date in the study of fatigue and motor learning has been on a performance decrement producing fatigue called reactive inhibition, which accumulates during continuous practice. It is generally held that with this reactive inhibition, the fatigue is more central in origin (i.e., analogous to boredom). Further, the work of Adams ( 1 , 2,), Digman (8, 9 ) and others (4, 5, 7) indicates that while performance is depressed by this central type of fatigue, | Fatigue is conceptualized as a *combined objective performance decrement* and *cognitive-resource depletion*. |

| | | | |
|---|---|---|---|
| | | learning itself is unaffected" (p. 682)<br><br>"A related problem which has not been as extensively investigated concerns the influence of physical fatigue - fatigue which is peripheral in origin (i.e., actual physical impairment) -upon motor learning." (p. 682) | |
| **COTTEN ET AL. 1974** | - | "While most evidence indicates that physical fatigue is detrimental to performance (1, 5, 6, 7, 8, 9, 13, 14, 16), the data on fatigue and learning is less conclusive." (p. 151) | Fatigue is conceptualized as an *objective performance decrement* |
| **COTTEN ET AL. 1972** | - | "The results (Alderman, 1965; Carron, 1969; Carron & Ferchuk, 1971; Godwin & Schmidt, 1971; Phillips, 1963; Schmidt, 1969; Stelmach, 1969) appear quite conclusive in indicating that performance is depressed by fatigue; however evidence concerning the effects of fatigue on learning is less clear." (p. 217) | Fatigue is conceptualized as an *objective performance decrement* |
| **DICKINSON ET AL. 1979** | - | "Richards (1968) postulated that pretask exercise generated both an exponential warm-up effect and a slower, but larger, exponential fatigue effect. The net result of these two components was to create an inverted-U relationship between amount of preliminary exercise and performance." (p. 81) | Fatigue is conceptualized as an *objective performance decrement* |
| **DWYER 1984** | - | "Depression of motor performance by physical fatigue is well established under a variety of experimental conditions. This autogenic fatigue may be analogous to reactive inhibition. In studies with fatigue generated from interpolated activities, performance decrements appear to be greatest when the motor task requires endurance and use of a large muscle mass."[…] (p. 130) | Fatigue is conceptualized as an *objective motor performance decrement* |
| **GODWIN AND SCHMIDT 1971** | - | "Since fatigue is well established as a variable which causes decrements in performance when | Fatigue is conceptualized as an |

| | | | |
|---|---|---|---|
| | | subjects are fatigued (i.e., it is a performance variable), fatigue should reduce the number of correct responses in a series of practice trials, and fatigued subjects should display less improvement with practice (on a subsequent transfer test under non-fatigued conditions) than relatively unfatigued subjects." (p. 374 – 375) | *objective performance decrement* |
| **HOSKENS ET AL. 2022** | **-** | "Cognitive fatigue potentially is also a method by which to suppress verbal working memory activity. Cognitive fatigue has been shown to reduce top-down conscious control processes (e.g., Borragán et al., 2016; van der Linden, 2011, 2003; Wolfgang & Schmitt, 2009). Wolfgang and Schmitt (2009), for example, found that prolonged performance of a Stroop task (480 trials) caused cognitive fatigue, which disrupted performance. […] Successful performance requires participants to consciously inhibit their automatic tendency to read and name the written word. Wolfgang and Schmitt (2009) argued that cognitive fatigue reduced cognitive resources available for top-down conscious inhibition of automatic responses (reading) during Stroop performance." (p. 1307) | Fatigue is conceptualized as a *depletion of cognitive resources* |
| **MASTERS ET AL. 2008** | **-** | - | |
| **MIERAU ET AL., 2009** | **-** | "In contrast, after high-intensity fatiguing exercise, both performance and/or learning has been found to be impaired [2,4,8–14]. Several authors have suggested an inverted U [15] effect of exercise intensity on motor performance and learning [1,8,11,16]. Thus, with increasing levels of exercise, performance should improve up to an optimal or maximal point and then decline again with a further increase in exercise intensity and/or duration." (p. 115) | Fatigue is conceptualized as an *objective performance decrement* |

| | | | |
|---|---|---|---|
| **NARDON ET AL., 2024** | | "Exercise-induced neuromuscular fatigue (NMF) is characterized by a temporary reduction in the ability of a muscle to produce force and/or power (1–3). This phenomenon affects various dimensions of human motor function, including the planning of motor activities (4–7), coordination (8–11), balance (12), sensorimotor integration (13, 14), limb proprioception (15, 16), and muscle activation patterns (17). Furthermore, NMF adversely affects gait characteristics (18–21) and posture stabilization (22–24). Specifically, NMF leads to postural instability by enhancing the sway velocity of the center of pressure (22) and diminishing the efficacy of balance recovery following disturbances (25)." (p. 629) | Fatigue is conceptualized as an *objective motor performance decrement* |
| **NUNNEY 1963** | Fatigue is considered a psychological phenomenon manifested in the subjective feelings of the individual, leading to a disinclination toward any form of work (no reference). (p. 370) | "**Fatigue is considered a psychological phenomenon manifested in the subjective feelings of the individual, leading to a disinclination toward any form of work.** On the other hand, the measurable physiological changes which are related to a loss of efficiency and effectiveness in body tissues are termed impairment" (p. 370) | Fatigue is conceptualized as *subjective feelings of tiredness/lack of energy* |
| **SIEKIRK ET AL. 2018** | - | "Fatigue is thought to reduce proprioceptive acuity and interfere with fluid movement (1, 4-6)." (p. 63)<br><br>"Some researchers have suggested that fatigue produces decrements in performance but may not have any effects on the learning of motor skills (9-12). Whereas others have reported that both performance and learning are impaired (4, 7, 13-17)." (p. 63) | Fatigue is conceptualized as a *combined objective performance decrement* and *cognitive-resource depletion* |
| **TAKAHASHI ET AL. 2006** | Fatigue is a commonly experienced condition that results from a period of intense | "**Fatigue is a commonly experienced condition that results from a period of intense or prolonged physical activity and is characterized by a reduced capacity to exert** | Fatigue is conceptualized as an *objective performance decrement* |

| | | | |
|---|---|---|---|
| | or prolonged physical activity and is characterized by a reduced capacity to exert muscular force (no reference) (p. 695) | **muscular force.** Fatigue is associated with mechanisms that are diverse and not completely understood but can develop due to factors proximal to the neuromuscular junction (20, 54, 61) or to factors involving the peripheral nervous system and muscle (1, 23, 35, 51). Fatigue slows muscle fiber conduction velocity (4, 43, 46), prolongs twitch duration (2, 18, 45, 64), and increases the neural activation required to produce a given force (8, 24, 26, 31, 63). Fatigue can affect motor performance in skilled activities such as targeted throwing (17), stoop lift (22), tennis (12), and balancing on an unstable surface (29)." (p. 695) | |
| **WHITLEY 1975** | - | "However, it should be noted that the findings concerning physical fatigue are somewhat conflicting. Fairly recent research (3, 6) shows that under certain conditions it can operate to significantly depress motor learning." (p. 111) | Fatigue is conceptualized as an *objective performance decrement* |
| **ZABIHHOSSEINIAN ET AL. 2020** | Fatigability (operationally defined as an inability to maintain task performance following an exercise intervention) (no reference) (p. 845) | "Cervical extensor muscle (CEM) **fatigability (operationally defined as an inability to maintain task performance following an exercise intervention)** has been shown to affect postural control, possibly due to central fatigue and/or proprioceptive conflicts (Gosselin et al. 2004)." (p. 845) | Fatigue is conceptualized as an *objective performance decrement* |

***STUDIES INVESTIGATING THE EFFECT OF FATIGUE INDUCED BY COGNITIVE EXERTION***

| | | | |
|---|---|---|---|
| **ANGUERA ET AL. 2012** | - | "The resource depletion framework offers a more direct way to probe the relationship between spatial working memory and visuomotor adaptation, given the view that higher cognitive processes are resource limited and can be temporarily depleted [cf. 14, 15–18]." (p. 107)<br><br>"Here, we rely upon a similar approach for Experiment 1: | Fatigue is conceptualized as a *combined objective performance decrement* and *cognitive-resource depletion* |

| | | | |
|---|---|---|---|
| | | selectively fatigue spatial working memory with intensive task performance of a spatial working memory task, and then evaluate performance on tasks that engage either related or unrelated cognitive processes." (p. 108) | |
| **APREUTESEI AND CRESSMAN 2024** | A psychobiological state caused by prolonged and/or intense periods of demanding cognitive activity and characterized by feelings of tiredness and lack of energy (Jacquet et al., 2021) (p. 2) | "[…] mental fatigue is defined as "**a psychobiological state caused by prolonged and/or intense periods of demanding cognitive activity and characterized by feelings of tiredness and lack of energy**" [19]. Altered states of consciousness and reduced alertness permeate our day to day lives." (p. 2) | Fatigue is conceptualized as *subjective feelings of tiredness/lack of energy* |
| **BANIHOSSEINI ET AL. 2025** | - | "The previous studies showed pervasive effect of physiological and mental fatigue on working memory, cognitive and motor performance that requires attention (Schmidt et al., 2018; Lam, 2008; Schmidt, 1975). Fatigue manifests as physical fatigue from prolonged tasks or mental fatigue stemming from extended cognitive engagement (Masters et al., 2008)." (p. 2)<br><br>"Mental fatigue has been shown to impair selective attention, influencing tactical performance in sports (Faber et al., 2012; Badin et al., 2016). Moreover, cognitive fatigue has been associated with enhanced procedural sequence learning by imposing stress on working memory, emphasizing the potential of implicit memory resources (Borragán et al., 2016; Yang et al., 2020)." (p. 2) | Fatigue is conceptualized as a *combined objective performance decrement* and *cognitive-resource depletion* |
| **BORRAGAN ET AL. 2016** | Decrease in cognitive resources developing over time on sustained cognitive demands independently of | "Another condition depleting the availability of controlled resources is mental or cognitive fatigue (CF), defined **as the decrease in cognitive resources developing over time on sustained cognitive demands independently of sleepiness** | Fatigue is conceptualized as a *depletion of cognitive resources* |

| | | | |
|---|---|---|---|
| | sleepiness (Trejoet al., 2005) (p. 2) | (Trejo et al., 2005). CF is associated with impaired cognitive control (Lorist et al., 2005), high-level information processing (Tanaka et al., 2012) and sustained attention (Langner et al., 2010).” (p. 2) | |
| **GODOI FILHO ET AL., 2025** | Psychobiological state characterized by tiredness, lack of energy, and apathetic feelings induced by long periods of demanding cognitive activity (p. 1) | “Mental fatigue is defined as a **psychobiological state characterized by tiredness, lack of energy, and apathetic feelings induced by long periods of demanding cognitive activity** (Smith et al., 2018; Van Cutsem et al., 2017). Evidence suggests that mental fatigue impairs physical performance, particularly endurance, strength, and power capacities (Van Cutsem et al., 2017).” (p. 1) | Fatigue is conceptualized as *subjective feelings of tiredness/lack of energy* and as an *objective performance decrement* |
| **KHOJASTEH MOGHANI ET AL. 2021** | Mental fatigue is a psychological state that results from constant cognitive activity; it is characterized by feelings of tiredness and lack of energy (Boksem &Tops, 2008; Marcora et al., 2009) (p. 2400) | “**Mental fatigue is a psychological state that results from constant cognitive activity; it is characterized by feelings of tiredness and lack of energy** (Boksem & Tops, 2008; Marcora et al., 2009). The negative effects of mental fatigue on cognitive tasks (Boksem et al., 2005, 2006) and motor tasks (Duncan et al., 2015; Habay et al., 2021; Magnuson et al., 2021) have been well demonstrated. Mental fatigue can damage the capability to maintain attentional focus (Boksem et al., 2005), monitor and adjust performance (Lorist et al., 2005), rapidly and accurately respond (Boksem et al., 2006), and identify and respond to important visual cues in action preparation (Boksem et al., 2006; Lorist et al., 2005).” (p. 2400) | Fatigue is conceptualized as *subjective feelings of tiredness/lack of energy* and as an *objective performance decrement* |
